# Photodegradation accelerates standing dead litter decomposition in monsoonal mountain grasslands of South America

**DOI:** 10.64898/2026.03.06.709891

**Authors:** Agustín Sarquis, Ignacio A. Siebenhart, Marcela S. Méndez, Amy T. Austin

**Author notes:** **Corresponding author:** Agustín Sarquis (email address).

## Abstract

Plant litter decomposition is the process through which plant-derived organic matter is recycled in terrestrial ecosystems. One of the drivers of decomposition is photodegradation, light-induced reactions that result in litter mass loss and transformations that accelerate the decomposition process. Photodegradation has been mostly studied in arid ecosystems, but mesic grasslands with monsoonal climate have been almost absent in the literature. In these ecosystems, standing dead biomass might remain exposed to solar radiation during dry winters while the effects of photodegradation accumulate. With the start of the warm and humid seasons, biotic decomposition might increase as a consequence of the changes in litter quality caused by sunlight. We aimed to study the impact of different wavelengths of solar radiation on litter mass loss and litter quality changes in a montane grassland with a monsoonal climate. We incubated litter from two dominant grasses under filters that generated treatments of full solar radiation, reduced UV radiation and reduced UV to short-wave visible radiation. We tracked changes in physical litter traits throughout the experiment under the three light treatment levels. We found an increase in litter mass loss due to sunlight exposure for both species, but each species reacted to a different range of wavelengths. We found evidence of enhancement of biotic decomposition by solar radiation (photofacilitation) in one of the two species, through an increase in *β*-glucosidase enzymatic activity. Seasonality affected litter decomposition of one species only by increasing mass loss depending on whether it was placed in the field during the dry winter or the humid spring. Finally, we found evidence of changes in physical litter traits caused by solar radiation, mainly in leaf mass per area (LMA) and water adsorption capacity. Our results represent the first proof of photodegradation in a productive grassland in this region, and highlight the fact that photodegradation is not constrained only to arid environments. Additionally, this study emphasizes the importance of litter physical traits in regulating carbon cycling through plant litter decomposition in terrestrial ecosystems.

**Open Research Statement:** Data are not yet provided as the study is currently under peer review. Upon acceptance, data will be archived in Zenodo. However, full datasets can be made available to the editorial board if required for the evaluation of the manuscript.

## Introduction

Plant litter decomposition is the process through which plant-derived organic matter is broken down into inorganic components. As a result, plant litter decomposition releases carbon (C) fixed by plants back to the atmosphere and intervenes in the formation of soil organic matter (SOM) (Cotrufo et al., 2015). The main biotic driver of decomposition is the metabolic activity of fungi and bacteria (Bradford et al., 2017), but soil fauna can also be important (Zanne et al., 2022; Lejoly et al., 2026). The magnitude of biotic decomposition is also determined by climate (Gholz et al., 2000) and litter chemistry (Cornwell et al., 2008; Wu et al., 2025; Zhang et al., 2008). Litter decomposition thus resides at a main intersection between C losses and sequestration in terrestrial ecosystems. For instance, rising atmospheric temperatures can increase CO_2_ emissions from litter decomposition (Hosseiniaghdam et al., 2023), reinforcing the greenhouse effect. Moreover, the formation of SOM from decomposing plant litter after decomposition is directly linked to soil fertility and soil C sequestration (Cotrufo and Lavallee, 2022). Hence, it is of great interest to reach a better understanding of how decomposition affects the terrestrial C balance and how this process might be affected by global change.

A lesser-known driver of litter decomposition in terrestrial ecosystems is photodegradation, the photochemical mineralization of organic matter due to exposure to solar radiation (Austin and Ballaŕe, 2024; King et al., 2012). It has been identified as important for its control on C release in a number of ecosystems, particularly in arid and semiarid zones (Austin and Vivanco, 2006; Berenstecher et al., 2020; Brandt et al., 2007; Day et al., 2007; Huang et al., 2017). Specifically, sunlight wavelengths in the range of ultraviolet (UV-B, 280-315 nm; and UV-A, 315-400 nm) and blue-green (400-550 nm) are largely responsible for these reactions (Austin and Ballaŕe, 2010; Brandt et al., 2009; Day and Bliss, 2019).

These photochemical reactions are possible due to the capacity of compounds in secondary cell walls to absorb sunlight, principally lignin, and produce photo-oxidative reactions (Austin and Ballaŕe, 2010; Kommedal et al., 2023; Moorhead and Callaghan, 1994). Moreover, this process is independent from microbial activity and can release C directly to the atmosphere as CO_2_, CH_4_ and CO (Brandt et al., 2010; Lee et al., 2012; Schade et al., 2012).

Photodegradation can also produce transformations in litter that make some carbohydrates to be more accessible to microbial consumption (Austin et al., 2016). This, in turn, increases litter mass loss as a complementary effect of sunlight called photofacilitation (Austin et al., 2016; Gallo et al., 2006; Méndez et al., 2022). For instance, exposure to solar radiation of litter can increase *β*-glucosidase enzymatic activity, an enzyme associated with the last step of cellulose degradation (Berenstecher et al., 2020, 2022; Méndez et al., 2019, 2022), and phenol-oxidase, associated with the degradation of phenolic compounds of lignin (Baker and Allison, 2015; Yao et al., 2022). It has been suggested that lignin degradation by sunlight could be responsible for this increase in biotic decomposition, due to the transformations of the lignin-cellulose matrix that increases microbial access to carbohydrates (Austin and Ballaŕe, 2024; Austin et al., 2016; Méndez et al., 2022). The importance of photodegradation was first appreciated in arid zones (Austin et al., 2006; Day et al., 2007; Gallo et al., 2009) and has been shown to be important in a variety of ecosystems, from deserts to arid grasslands and woodlands (Almagro et al., 2015, 2017; Baker and Allison, 2015; Brandt et al., 2010; Day et al., 2018; Henry et al., 2008). These arid and semiarid ecosystems are characterized by low plant productivity and heterogeneous vegetation, which contribute to the interception of solar radiation by plant litter. Photodegradation has been evaluated in both monsoonal (Brandt et al., 2010; Day et al., 2022; Pancotto et al., 2003) as well as Mediterranean climates (Austin et al., 2006; Ruhland and Fraley, 2023; Rutledge et al., 2010) of low mean annual precipitation (MAP). These studies have proven that sunlight affects decomposition in both types of climates, with divergent patterns at local scales. In Mediterranean climates, the peak of solar radiation in summer does not coincide with the rainy season, whereas in monsoonal climates peaks in solar radiation and precipitation coincide. This suggests that whether peaks in radiation and rains are synchronous or not can determine how abiotic and biotic factors influencing decomposition may interact with each other. More recent research on photodegradation has expanded to other types of ecosystems including temperate forests, tropical ecosystems and agroecosystems (Keiser et al., 2021; Marinho et al., 2020; Wang et al., 2021; Cabrera et al., 2026), but the majority of mesic ecosystems have not yet been evaluated for the potential importance of solar radiation on C turnover, particularly for the relative importance of direct photodegradation and photofacilitation effects (Austin and Ballaŕe, 2024).

Considering grasslands are some of the most productive and widespread ecosystems on Earth (Gibson, 2008), it would be important to advance our understanding of how solar radiation affects this portion of the terrestrial C balance (Keiser and Nieland, 2025). Currently, most studies in grasslands have been set in low-rainfall sites, many of which showed marked positive effects of photodegradation (Yang et al., 2025; Butler et al., 2023; Wang et al., 2017), but some of them found either neutral or negative effects as well (Erdenebileg et al., 2018; Yang et al., 2024; Almagro et al., 2017). However, fewer studies have been carried out in more productive grasslands with higher mean annual precipitation (MAP) (Brandt et al., 2010; Butler et al., 2023; van Asperen et al., 2015). These grasslands are particularly relevant for the question of the importance on photodegradation due to the accumulation of very large quantities of standing dead material associated with tussock grass formation, also known as marcescence (Mudŕak et al., 2023; Sarmiento, 1992). These large amounts of plant litter are exposed to solar radiation for long periods before touching the ground. Circling back to seasonality in these grasslands, even fewer studies have been done in grasslands with monsoonal climate (Yao et al., 2024). It is worth exploring whether in monsoonal grasslands abiotic photodegradation acts during the dry winter and photofacilitation begins at the onset of the humid and warm seasons.

Several field studies have also specifically addressed the effect of litter traits on photodegradation and photofacilitation. This is important because part of how sunlight impacts decomposition is through modifying litter chemistry and structure and thus changing its decomposability (Austin et al., 2016; Foereid et al., 2010; Gallo et al., 2006). Most studies have focused on chemical traits like C, N and C fractions like lignin, hemicellulose and cellulose, (Ball et al., 2019; Day et al., 2022; Ma et al., 2017; Yang et al., 2025). The focus on these traits, specifically on lignin, is because of its importance as a control on both biotic decomposition and photodegradation (Austin and Ballaŕe, 2010). However, there is a need to expand studies to incorporate other litter traits that might potentially be linked to photodegradation, including how those traits change over time. In that regard, few physical traits have been measured in photodegradation studies in the field, typically SLA/LMA (Wang et al., 2024; Li et al., 2024; Gaxiola and Armesto, 2015), leaf toughness (Araujo et al., 2022; Masubelele and Bond, 2022), water and vapor holding capacity (Almagro et al., 2017; Logan et al., 2022), and light-interacting traits (i.e.: reflectance, transmittance, and absorptance) (Day and Bliss, 2020; Day et al., 2015; Ruhland and Fraley, 2023). There is a clear opportunity for the advancement of photodegradation research in how physical litter traits change over time and how this feeds back to C release at the ecosystem scale.

Our objectives in this study were to understand the importance of photodegradation for C turnover and to evaluate the spectral dependence on C release from plant litter in a monsoonal mountain grassland in central Argentina. Previous studies for the region were carried out in grass-dominated ecosystems with similar MAP and mean annual temperature (MAT), but with a different rainfall seasonality (i.e. Mediterranean climate; Berenstecher et al., 2020; Méndez et al., 2019). We specifically designed our study to directly address the seasonal contributions and the role of dominant species identity in the relative importance of sunlight on litter decomposition of standing dead biomass. We hypothesized that exposure to sunlight accelerates C cycling throughout the year in this monsoonal grassland, and that this relationship is modulated by seasonality. During dry winters, sunlight acts predominantly through the abiotic photo-oxidative route. Instead, during the warm and humid seasons, direct abiotic photodegradation and biotic decomposition occur simultaneously. Additionally, sunlight exposure during the dry season promotes biotic decomposition during the following humid season through changes in litter quality (photofacilitation). Lastly, photodegradation of the litter produces changes in its physical quality over time. We applied a temporally asynchronous design to disentangle the relative importance of contrasting seasons in order to test our hypothesis. Only a few studies have tried to address directly how marked seasonal differences affect litter decomposition with this design (i.e.: Berenstecher et al., 2020; Li et al., 2024). We found that both seasonality and species identity and their associated litter traits were key in defining the importance of photodegradation as a control on decomposition in this montane grassland.

## Methods

### Study site

Our study site was located in a native montane grassland in a plateau known as Pampa de Achala located in the Córdoba mountain ranges of central Argentina (2000 – 2300 m a.s.l.). The mean temperatures of the coldest and warmest months are 5.0 and 11.4 °C, respectively. Average mean annual precipitation is 900 mm, highly concentrated between October and April, and frosts occur year round (Cingolani et al., 2015). Additionally, fog events are common year round and represent an additional water input into the system (Poca et al., 2018). Soils are Mollisols derived from granitic rocks and fine texture particles originated from wind erosion (Cabido et al., 1987). The landscape is an undulating plain with patches of rocky outcrops, *Polylepis australis* Bitter woodlands, and short-statured grazing lawns, all immersed in a tussock grassland matrix with varying degrees of openness dominated by *Poa stuckertii* (Hack.) Parodi and *Deyeuxia hieronymi* (Hack.) Türpe (Cingolani et al., 2014; von Müller et al., 2017; Zeballos et al., 2024). These tussock grassland physiognomies are highly productive and generate each year large quantities of senescent material during the dry season (fall-winter) that stay attached to the plants for extended periods (Pucheta et al., 1998). This implies that a considerable amount of dry biomass is exposed to sunlight during an extended period of time, which makes this system an ideal laboratory to study the effects of photodegradation. In this system, the relatively high stocking rates and wildfires help maintain the grazing lawns (Cingolani et al., 2013), while in zones with lower stocking rates, tall tussock grasses dominate (i.e. *P. stuckertii* and *D. hieronymi*). We worked within the limits of Quebrada del Condorito National Park, where grazing is used as a management tool with conservation aims and to stop the accumulation of excessive amounts of flammable standing dead biomass (Cingolani et al., 2014; von Müller et al., 2017). The rangeland where we carried out this study had an effective stocking rate for the period 2020-2021 of 0.13 AU *ha^−^*^1^ (Animal Units per hectare) (Administración de Parques Nacionales, 2021).

### Experimental design

The general experimental design consisted of the decomposition in the field of two dominant native grass species, *P. stuckertii* and *D. hieronymi*. Leaf litter was decomposed in plots covered in the respective species, hanging as if it were standing dead biomass. Each species was subjected to three levels of a sunlight treatment to evaluate the importance of photodegradation as a driver of litter decomposition in this site. We deployed samples in the field at three different dates to study the impact of seasonality on decomposition, and we collected the samples from the field on four dates along a 2-year period.

In more detail, we established 5 pairs of plots on the same hillside with grassland cover (Figure 1a) for the evaluation of photodegradation and seasonality on native plant litter decomposition (31a 37’ S, 64a 48’ W; 2150 m a.s.l.). Out of each pair of plots one had *P. stuckertii* cover and another one, *D. hieronymi* cover. Samples of each of the two species were later incubated in plots covered in their respective species. Plots were located south of rocky pavements to avoid shading from vegetation. We chose this exposure for the plots to maximize sunlight exposure in the Southern Hemisphere.

**Figure 1:**
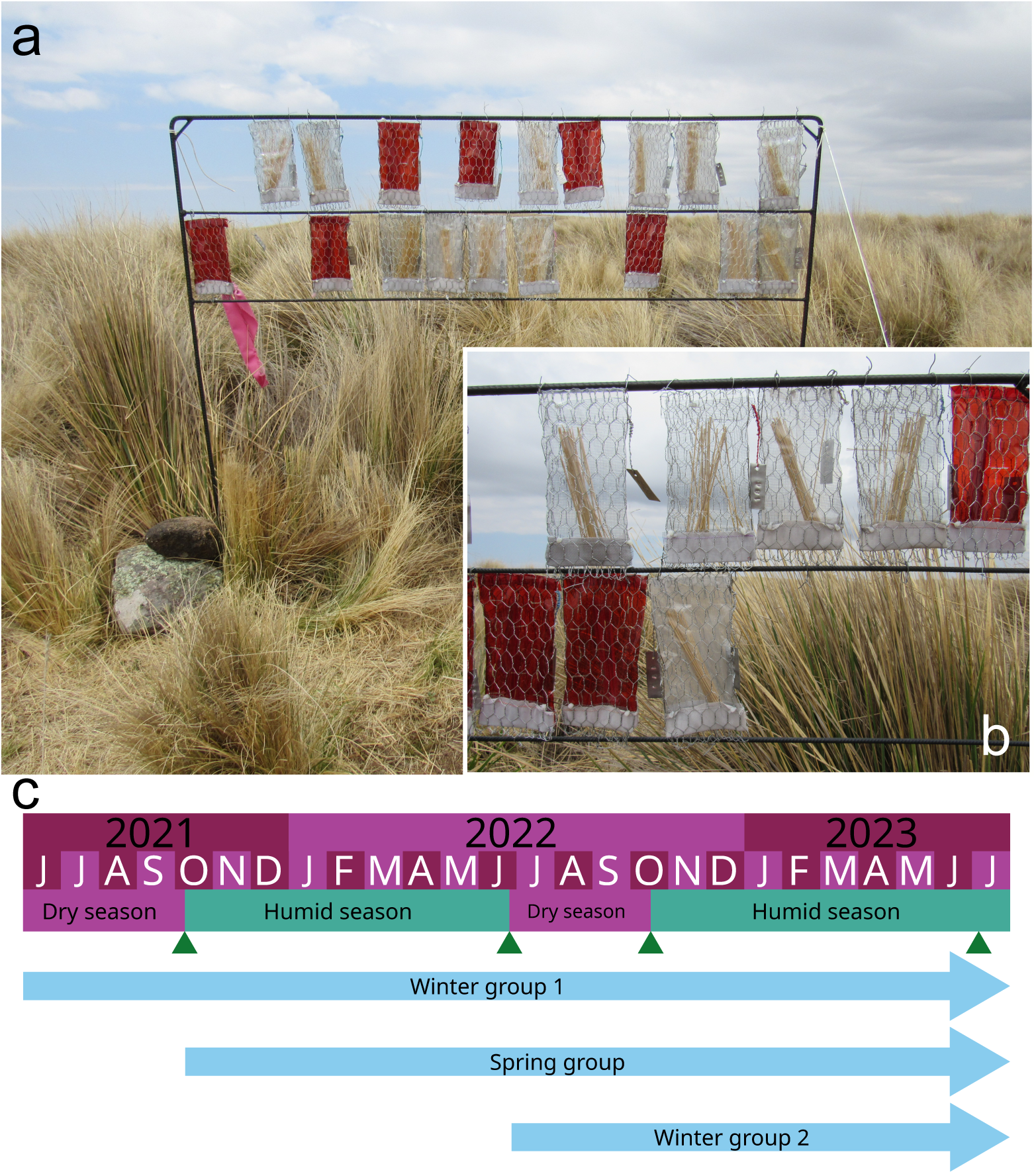
Field plot with *D. hieronymi* cover (a); close-up of litter envelopes of *P. stuckertii* (b); time frame for the litterbag placement and overall experimental design (c). Dark green triangles mark sample collection dates. Photographs by A. Sarquis.

In July 2019, we collected standing dead material (hereafter ‘litter’) of *P. stuckertii* and *D. hieronymi* in the same rangeland where the experiment was done, and in a neighboring rangeland. We took the litter to Buenos Aires where we stored it in boxes in a dry and cool space in the Instituto de Investigaciones Fisiológicas y Ecológicas Vinculadas a la Agricultura (IFEVA). We made sure to flip the litter in the boxes periodically to keep it aired until processing. We selected senescent blades with the least signs of decomposition. We cut blades in 10-16 cm segments and weighed samples of 1.5 g of air-dried litter. We preserved 10 samples for measurements of initial litter quality, and the rest were randomly assigned to a treatment.

To evaluate mass loss, we designed envelopes made of hexagonal galvanized wire mesh with opening of 13 mm, 20 cm in height and 10 cm in width (Figure 1b). We oriented the envelopes on the hangers in a north-south direction. The north side of the envelopes had a plastic filter that blocked a specific range of the wavelength in order to create three distinct sunlight treatment levels: full radiation (R+), blocked ultraviolet (UV-), and all photochemically active radiation blocked (R-). The R+ filter was a 100 *µ*m thickness polyethylene with a transmittance from UV-B to the visible spectrum (Appendix S1: Figure S1; 280-800 nm; Ever Wear S. A.). The UV-filter blocked radiation in the UV range (280-400 nm; Costech, 226 UV). The R-filter blocked radiation from UV to blue-green light (280-550 nm; Rosco@ Na135 Deep Golden Amber). This type of filter has proven to be efficient in reducing radiation with photo-oxidative effects in several other studies (Austin and Ballaŕe, 2010; Brandt et al., 2009; Day and Bliss, 2019). It is worth noting that these filters cannot separate the abiotic effects of photodegradation (direct photomineralization) from the biotic effects (photofacilitation), nor the deleterious effects of sunlight on microbes. Thus, these filters allow us to demonstrate an overall sunlight effect that is the balance of these three processes. We punched holes in the filters and covered the southern side of the envelopes with fiberglass mesh of 2 mm openings to allow the passage of air and humidity. We added a polyester gauze pocket to the bottom of the envelopes to avoid fragmentation and loss from the bottom.

Back in the field, within each plot we constructed a horizontal iron hanger 1 m high and 1.6 m long in front of the vegetation. We hung the envelopes with litter samples in two rows at 60-80 and 80-100 cm high. These heights are within range for the vegetation in the area (Vaieretti et al., 2010), thus we simulated decomposition of standing dead material. We measured temperature of the samples in the field on 5 occasions using an infrared thermometer (model 63, Fluke), and we did not find a significant effect of filters on litter temperature (Appendix S1: Table S1).

Samples were placed and collected along a two year trajectory in order to evaluate the effect of different seasons on photodegradation (Figure 1c). The first group of samples began in June 2021, at the onset of the dry and cold season and it lasted for 2 years on the field (hereafter Winter Group 1). The second group started on October 2021, at the onset of the warm and humid season, and it lasted for 1.7 years (hereafter Spring Group). The last group started on June 2022, and it lasted for 1 year (hereafter Winter Group 2). We retrieved samples on 4 dates at 0.3, 1, 1.3 and 2 years. Each time we collected one litter envelope per group, species and light treatment per plot. In total we had 270 samples: 2 species x 5 plots x 3 sunlight levels x (4 harvests for Winter Group 1 + 3 harvests for Spring Group + 2 harvests for Winter Group 2). On each collection date, we put samples in paper envelopes and sealed them in plastic zipper bags individually for their transportation to Buenos Aires. We kept samples cold with ice packs during transportation and stored them in a freezer until processing.

### Mass loss

First, we brushed samples and cleared extraneous material with tweezers. We measured fresh mass and extracted 0.200 g for enzymatic activity assays (see below). We dried the rest of the sample at 50 °C for 48 h and measured dry mass. We calculated water content and used it to estimate the dry mass of the sub-sample used for enzymatic activity assays and finally calculated total sample dry mass. We ground samples and corrected for ash content using combustion at 450 °C in a muffle furnace (Harmon et al., 1999). We fit a negative exponential model for each plot using lineal regression following equation

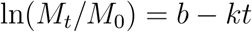

where *M*_0_ is ash-free initial dry mass, *M_t_* is ash-free dry mass at time *t*, *b* is the intercept, and *k* is the slope, also called decomposition rate (Olson, 1963). We only used *k* constants at the plot level if the regression was significant (*p <* 0.05). We did not use data from Winter Group 2 because it was not possible to fit the model with confidence for only 3 time points. Because of that, for samples that stayed in the field for a year we calculated daily organic mass loss as an alternative. In this way, we compared daily mass loss up to 1 year and *k* until 2 years.

### Physical litter traits

We measured leaf area of initial and field samples, except for the October 2021 date. We used a scanner (model V370, Epson, Japan) and analyzed images with ImageJ software (Schneider et al., 2012). We calculated leaf mass per area (LMA, mg mm*^−^*^2^) dividing dry mass by area (Pérez-Harguindeguy et al., 2013). We measured leaf toughness using the punch test (Pérez-Harguindeguy et al., 2013), for initial samples, and samples from Winter Group 2 at the October 2022 date, and for all groups at the July 2023 sampling date. We used a digital dynamometer (DFX2-010, Chatillon) coupled with a flat needle 1.91 mm in diameter. To calculate force to punch, we divided the force by the perimeter of the needle (N mm*^−^*^1^). When blades were thinner than the needle, we approximated the perimeter to a rectangle long as the diameter of the needle, and wide as the sample. Blades of *P. stuckertii* and *D. hieronymi* fold at senescence, thus it was necessary to unfold them before measuring. However, this was not possible for *D. hieronymi* because it is too fine and fragile when dry. So, *D. hieronymi* values correspond to the force to punch two tissue layers. We recognize this is not comparable to other measurements, but it is more realistic given that *D. hieronymi* litter persists in this form during decomposition.

We measured water adsorption capacity with a technique modified from Day et al. (2022). This is a measure of hydrophilicity of litter. We did this for initial samples, for Winter Group 2 at the October 2022 date, and for all July 2023 samples. We dried samples of 0.250 g at 60 °C for 48 h and registered initial dry mass (*W_i_*). We put samples in nylon bags (40 µm pores) and submerged them in distilled water for 30 min. After draining the bags, we dried excess water adhered to samples with paper and measured wet mass (*W_w_*). We dried samples again at 60 °C for 48 h and measured final dry mass (*W_f_*). We calculated water adsorption capacity as (*W_w_ − W_f_*)*/W_f_*, as percentage. We also calculated potential leaching mass loss as (*W_i_ − W_f_*)*/W_i_*, as percentage.

### Chemical litter traits

We measured initial soluble carbohydrates, hemicellulose, cellulose and lignin with the acid detergent sequential digestion method (Van Soest et al., 1991). We put 0.5000 g (*w*_0_) of dry ground samples in Ankom F57 filter bags and put them in a sequential automatized analyzer (Ankom 220, Ankom Technology, NY, USA). We first washed samples in water at 25 °C for 1 h, dried them for 24 h at 100 ° C and measured dry mass (*w*_1_). Then, we performed an acid detergent digestion at 100 °C for 1 h, after which we washed samples 3 times in water at 90 °C and twice in acetone, for 15 min each time. We dried samples and measured dry mass (*w*_2_). After this, we put samples in sulfuric acid at a 72% concentration for 3 h, washed 3 times with hot water and twice with acetone. We dried samples and measured dry mass (*w*_3_). We finally combusted samples in a muffle furnace at 450 °C for 4 h and calculated ash content (*w*_4_) to correct for inorganic mass. We calculated the proportion of soluble carbohydrates as (*w*_0_ *− w*_1_)*/*(*w*_0_ *− w*_4_), hemicellulose as (*w*_1_ *− w*_2_)*/*(*w*_0_ *− w*_4_), cellulose as (*w*_2_ *− w*_3_)*/*(*w*_0_ *− w*_4_) and lignin as (*w*_3_ *− w*_4_)*/*(*w*_0_ *− w*_4_). We measured total initial sugars following de DuBois et al. (1956). First, we digested 0.035 g of dry grinded samples in 7 ml of HCl 2.5 N at 100 °C for 3 h. We neutralized it with NaCO_3_ and added 93 ml of distilled water. We took an aliquot of 1 ml and added 1 ml of 5% phenol and 5 ml of 96% sulfuric acid. We measured absorptance at 490 nm with a UV-VIS spectrophotometer (Shimadzu Scientific Instruments, Japan). We calculated total sugar concentrations with a calibration curve made using a solution of dextrose in distilled water.

We measured initial saccharification (accessibility to cell wall polysaccharides by microbial enzymes) (Breuil and Saddler, 1985; Chen and Dixon, 2007; Ghose, 1987; Méndez et al., 2022). We prepared a 50 U/ml solution of the *Trichoderma viride* cellulase enzyme (C9422, Sigma Aldrich) in acetate buffer 50 mM at 5.5 pH. We added 1 ml of enzymatic solution, 4 ml of buffer and 0.2 ml of toluene to tubes with 0.0250 g of dry grinded samples. We incubated the tubes for 72 h at 50 °C under constant shaking. We took 1 ml aliquots and added 2 ml of distilled water and 3 ml of a solution 100 ml solution made from 1% NaOH with 1 g of dinitrosalicylic acid, 0.2 g of sodium sulfite and 0.05 g of phenol. We incubated the samples for 5 min at 100 °C, after which we added 1 ml of Rochelle salt solution at 40% (40 g of sodium and potassium tartrate in 100 ml of distilled water). Once the samples were chilled, we measured absorptance at 575 nm. We quantified sugar concentrations using a calibration curve.

We extracted total polyphenols from 40 mg of dried ground sample in 20 mL of 50% methanol at 80 °C for 1 h. We determined polyphenols concentration with the Folin-Ciocalteu method (Cadisch and Giller, 1997), measuring absorptance at 760 nm. For the quantification we used a gallic acid calibration curve. We also measured sunscreens (hereafter A305), which are phenolic compounds that absorb light at 305 nm (Mazza et al., 2000). We extracted A305 with methanol:HCl 99:1 for 48 h at -20 °C and measured absorptance at 305 nm.

### Enzymatic activity

We measured potential *β*-glucosidase and phenol-oxidase enzymatic activities of litter incubated in the field at each date (Sinsabaugh et al., 1999). We put 0.200 g of fresh litter in 7 ml of distilled water and shook it. For *β*-glucosidase activity we prepared a 10 mM solution of p-nitrophenyl-*β*-D-glucopyranoside substrate in 50 mM acetate buffer at 5.5 pH. For each sample we prepared a tube with a 1 ml of litter solution plus 1 ml of substrate, and a tube with 1 ml of litter solution plus 1 ml of buffer. We also prepared tubes with 1 ml of substrate and 1 ml of distilled water as blank tubes. We incubated the tubes for 2 h at 24 °C and centrifuged them. We stopped the reaction with 0.2 ml of 1M NaOH and measured absorptance at 410 nm in a spectrophotometer.

To measure phenol-oxidase enzymatic activity, we prepared a 12.5 mM solution of the 3,4-dihydroxy-L-phenylalanine substrate with the same acetate buffer as before. For each sample we prepared a tube with 2 ml of litter solution plus 1 ml of substrate, and a tube with 2 ml of litter solution plus 1 ml of buffer. We also prepared blank tubes with 1 ml of substrate and 2 ml of distilled water. We incubated the test tubes for 3 h at 24 °C and centrifuged them. We measured absorptance at 460 nm with a spectrophotometer. For both enzymes we calculated their activity (*A*) as

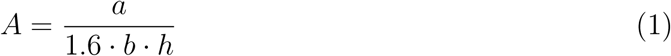

where *a* is net absorptance, 1.6 is the extinction coefficient in *µ*M, *b* is the litter sample in g per ml of aliquot, and *h* is incubation time in hours.

### Statistical analysis

We used ANOVA to analyze remaining organic matter and accumulated enzymatic activity comparing light filters per each date, species and starting group separately. We also analyzed remaining organic matter between June and October 2022 dates per filter and species. We only compared these dates because we were interested in knowing what happened during the dry cold season. We compared daily mass loss and *k* constants between filters per species and starting group separately using ANOVA. We also compared these variables between starting groups per species only for filter level R+ as a way to compare the impact of seasonality on decomposition under near-ambient conditions. We analysed differences in initial traits between the two species with ANOVA. We removed one value leaf toughness and A305 that was defined as an outlier based on Cook’s distance.

We tested normality of errors using Shapiro-Wilk’s test and homoscedasticity with Levene’s test. To fulfil ANOVA assumptions, we transformed using natural logarithm accumulated *β*-glucosidase activity results from *P. stuckertii* of Winter Group 1 at 1 year, and from Winter Group 2 in all its dates. We transformed using ln +1 results from accumulated phenol-oxidase activity of Winter Group 1 at 1.3 and 2 years, and of Spring Group at 0.7 years. We did not include anomalous data from plot 6 of phenol-oxidase activity of *P. stuckertii* of Spring Group at 1 year. We could not fulfil normality assumption of phenol-oxidase activity in *P. stuckertii* of Winter Group 1 at 1 year because of excess of zeros, and using mixed models for zero-inflation did not improve model fit. Finally, to evaluate changes in litter traits over time in comparison with initial values, we performed t-tests and calculated 95% confidence intervals. We performed all statistical analyses in R (R Core Team, 2020).

## Results

### Effects of sunlight and season on litter decomposition and enzymatic activity

Results for organic mass loss of litter and accumulated *β*-glucosidase activity during the two years of the experiment are shown in Figure 2 only for Winter Group 1 (ANOVA results in Appendix S1: Table S2). After 1.3 years of field exposure, *P. stuckertii* showed a 15% significant increase in mass loss due to visible light (Figure 2a). Interestingly, decomposition seemed to halt during the dry (winter) season, since there were no detectable differences between June and October 2022 for any attenuation treatment (Figure 3; *p*: 0.2, *R*^2^: 0.3). *D. hieronymi*, in turn, showed a relative increase of 46% in mass loss due to UV light at 1.3 years (Figure 2b). Here, there was also a halt in decomposition during the dry season for UV-(*p*: 0.3, *R*^2^: 0.2) and R-(p: 0.7, *R*^2^: 0.02) filters, but not for full sunlight filters (R+) which lost 5.0 ± 2.1 % more mass during that period (Figure 3; *p*: 0.047, *R*^2^: 0.5). After 2 years in the field, UV light increased mass loss by 30% (Figure 2b). Accumulated *β*-glucosidase activity in *P. stuckertii* increased over time but showed no clear response to solar radiation attenuation (Figure 2c). In contrast, *D. hieronymi* presented a significant 50% increase in enzyme activity after 2 years in the field due to UV exposure (Figure 2d). Phenol-oxidase activity did not have a clear response to sunlight in this experiment (Appendix S1: Table S3). Full results for Spring Group and Winter Group 2 can be found in the Appendix S1 (Figure S2, Table S4). Generally, results from the other two groups followed similar seasonal patterns in response to water availability compared to Winter Group 1.

**Figure 2:**
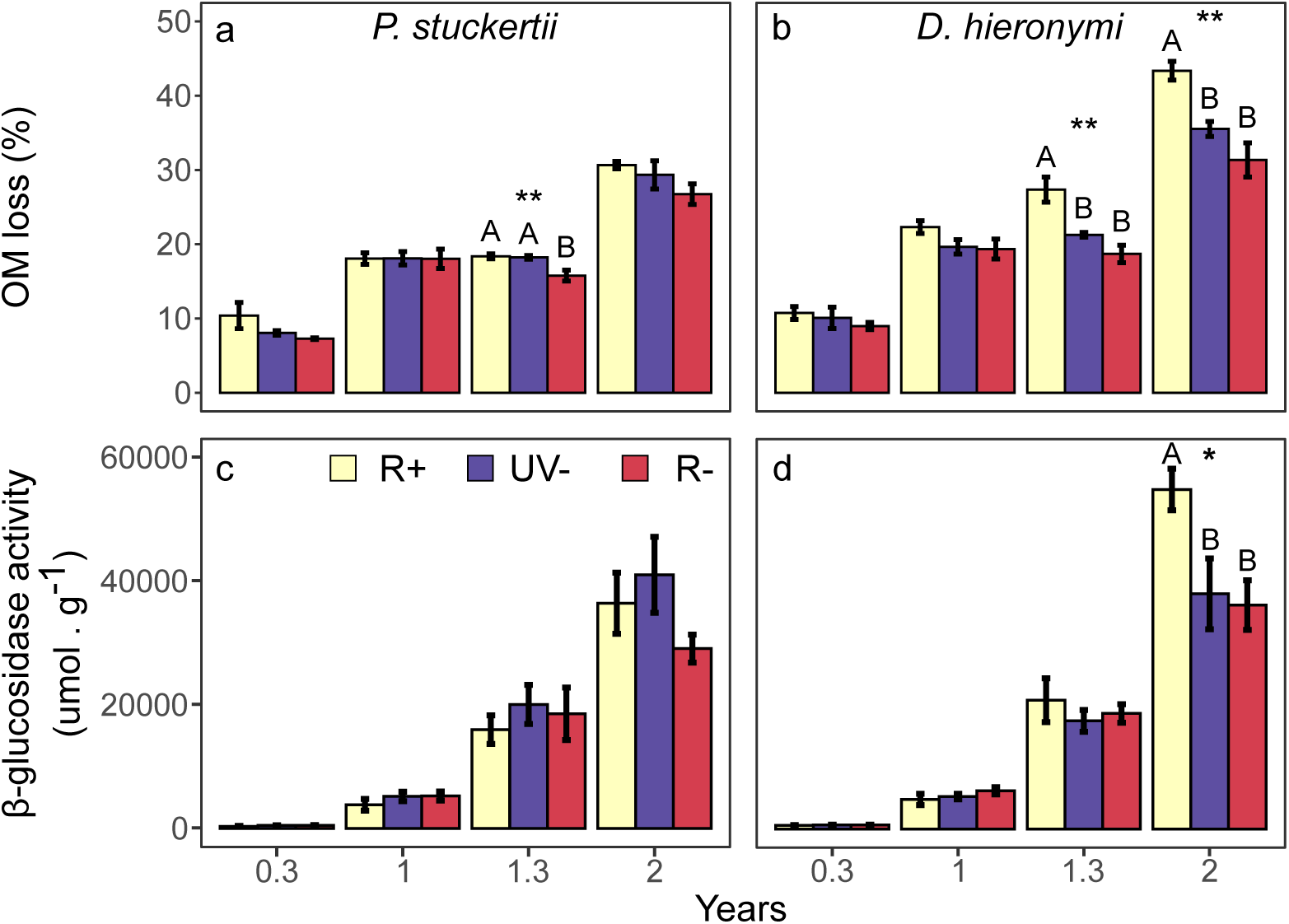
Organic mass (OM) loss (%; a, b) and accumulated *β*-glucosidase activity (*µ*mol g*^−^*^1^; c, d) over time for both species of Winter Group 1. Asterisks denote significant differences between filters per date: ** *p <* 0.01, * *p <* 0.05. Upper-case letters denote differences between filters following Tukey HSD test. Bars represent mean values and standard errors. Results in this figure show OM mass loss, but statistical analyses were performed on remaining OM. R+: full radiation. UV-: UV blocked. R-: UV, blue and green light blocked.

**Figure 3:**
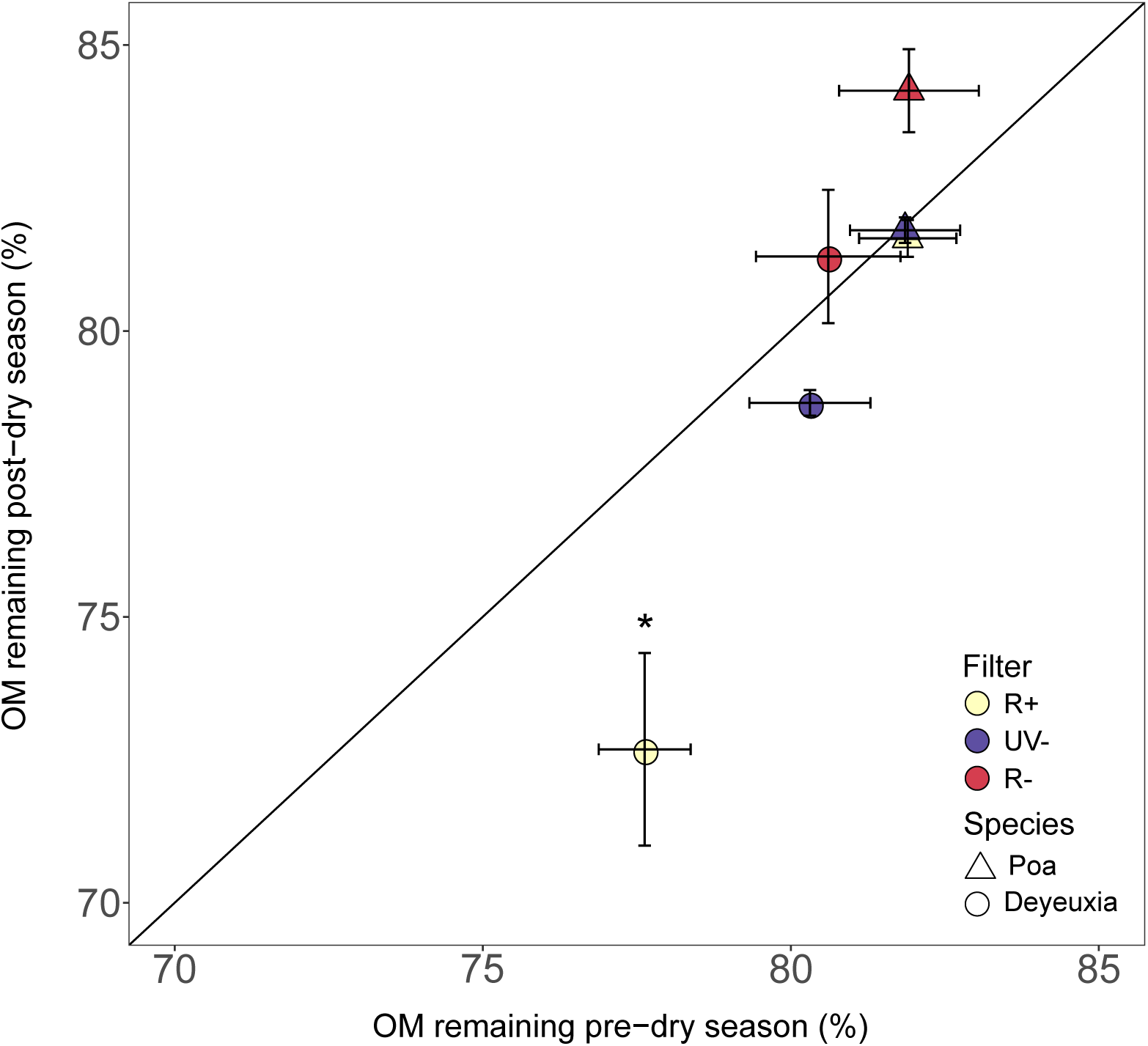
Organic mass (OM) remaining (%) pre- and post-dry season of 2022. Shapes are mean values and bars are standard errors. Asterisk denotes significant differences between dates per filter: * *p <* 0.05. Only Winter Group 1 values are included in this figure. R+: full radiation. UV-: UV blocked. R-: UV, blue and green light blocked.

### Litter decomposition across seasons and dominant grassland species

Decomposition for litter starting in winter (Winter group 1) and in spring are shown in Figure 4 (ANOVA results in Appendix S1: Table S5). For results after 1 year in the field, we show daily mass losses. *P. stuckertii* did not show a response to light filters in daily mass loss at 1 year for any of the seasonal groups (Figure 4a). There was however a significant effect of starting season, since R+ samples decomposed 18% faster when they were placed to decompose in winter, when compared to spring (*p*: 0.01, *R*^2^: 0.6). For results at the end of the experiment, we show *k* values (yr*^−^*^1^). Radiation attenuation had a significant effect on integrated decomposition, with a 19% increase at the spring start. Light treatments did not differ significantly between filters for the winter group for *P. stuckertii* (Figure 4c). Additionally, the effect of starting season at the end of the experiment switched compared to 1 year results. R+ samples that started decomposing in the spring decomposed 23% faster compared to those that started in the winter (*p*: 0.02, *R*^2^: 0.5).

**Figure 4:**
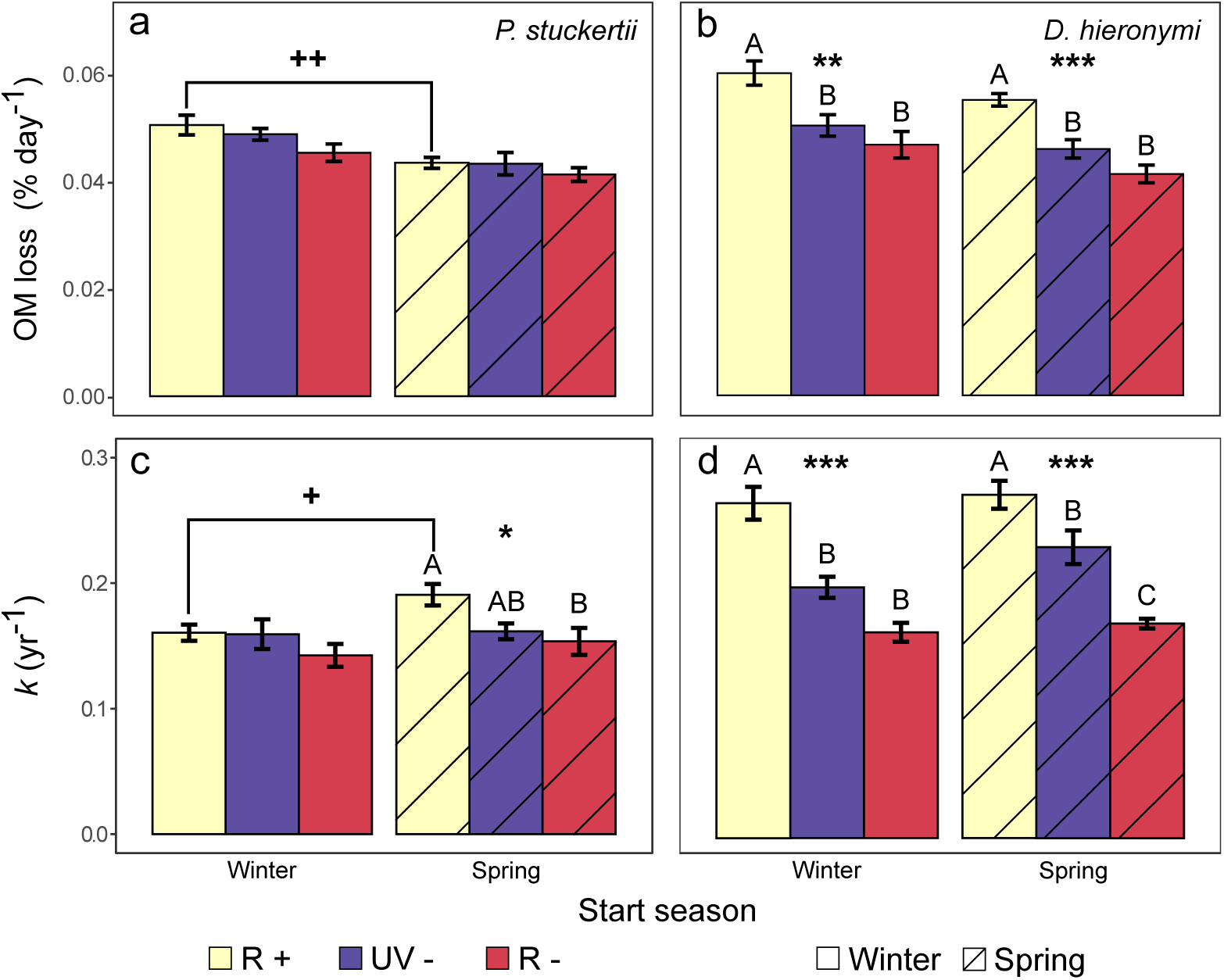
Decomposition rates for each filter by species and starting group. Daily organic mass (OM) loss (% day*^−^*^1^; a, b) was calculated for samples after 1 year and *k* constants were calculated at the end of the experiment. Asterisks denote significant differences between filters of the same starting group: *** *p <* 0.001, ** *p <* 0.01, * *p <* 0.05. Upper-case letters denote differences from Tukey test between filters of the same starting group. Plus signs denote significant differences between starting groups (for filter level R+ only): ++ *p <* 0.01, + *p <* 0.05. Bars are mean values with standard errors. R+: full radiation. UV-: UV blocked. R-: UV, blue and green light blocked.

In the case of *D. hieronymi*, daily mass loss at 1 year increased due to UV exposure by 25% and 20% for the spring and winter groups, respectively (Figure 4b). In contrast, the season of start did not influence daily mass loss for this species (*p*: 0.08, *R*^2^: 0.4). At the end of the experiment, *k* values of the winter group showed a 56% increase due to UV light (Figure 4d). Instead, the Spring Group reacted to both UV and visible light with a 24% and 35% increase, respectively. Again, this species did not react to season of start (*p*: 0.7, *R*^2^: 0.02). Overall, decomposition rates suggest that *P. stuckertii* responds more to seasonality, while *D. hieronymi* responds more to sunlight exposure.

### Initial litter quality

We observed differences in physical and chemical initial litter traits between both species (Figure 5, Appendix S1: Table S6). Out of 7 chemical traits, only 3 significantly differed between species. *P. stuckertii* litter had 22% less absorptance at 305 nm, 15% more cellulose, and 8% less hemicellulose than *D. hieronymi*. Additionally, all 4 physical traits differed between species. *P. stuckertii* litter was almost three times as tougher, twice as denser (higher LMA), adsorbed a third more water, and lost almost 7 times more mass through leaching than *D. hieronymi*.

**Figure 5:**
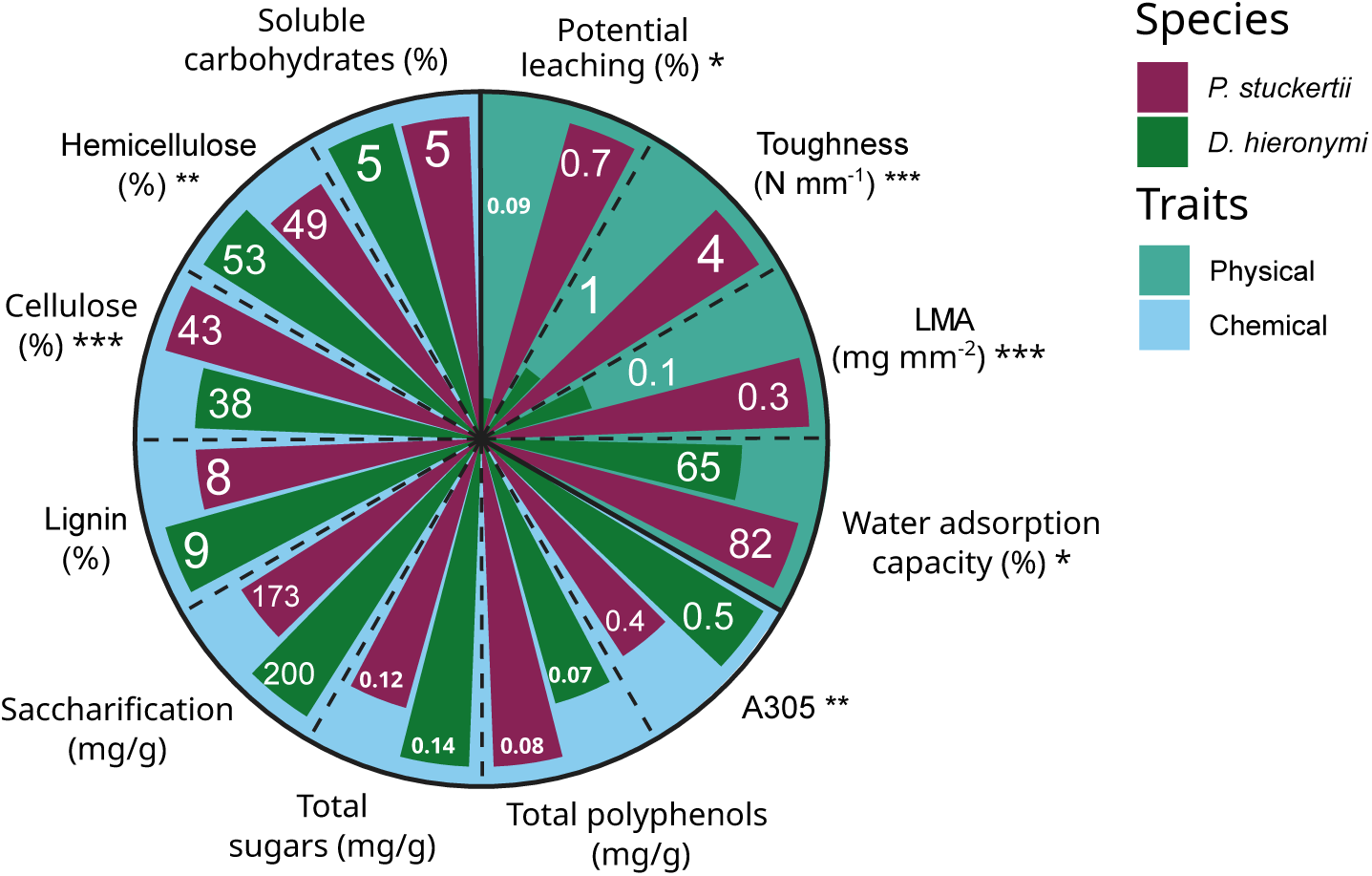
Initial physical and chemical initial traits of the two species. Asterisks denote significant differences between species: *** *p <* 0.001, ** *p <* 0.01, * *p <* 0.05. A305: sunscreens with absorptance at 305 nm. LMA: leaf mass per area.

### Changes in litter traits

There were substantial changes in physical litter traits during the field incubation with respect to initial values (Figure 6). Starting with *P. stuckertii*, its LMA decreased around 9% as a response to visible light at 0.7 years (*p*: 0.009). This pattern continued until 1.7 years when litter under all filters had lower LMA than initially. Water adsorption capacity increased by 33-41% at the 1.7 year sampling date due to exposure to visible light with respect to initial values (*p*: 0.0008). At the end of the experiment, however, all samples became more hydrophilic (R+ samples too, even when the effect was not significant, the mean relative effect had the same magnitude as the other treatment levels). Leaf toughness in *P. stuckertii* showed an apparent hardening at 0.3 years (only significant under R-; *p*: 0.04), but overall, leaf toughness decreased for all samples over time independently of light treatment. Finally, there was initially an increase in potential leaching (mass lost after soaking in water), followed by a decrease until the end of the experiment (Appendix S1: Figure S3).

**Figure 6:**
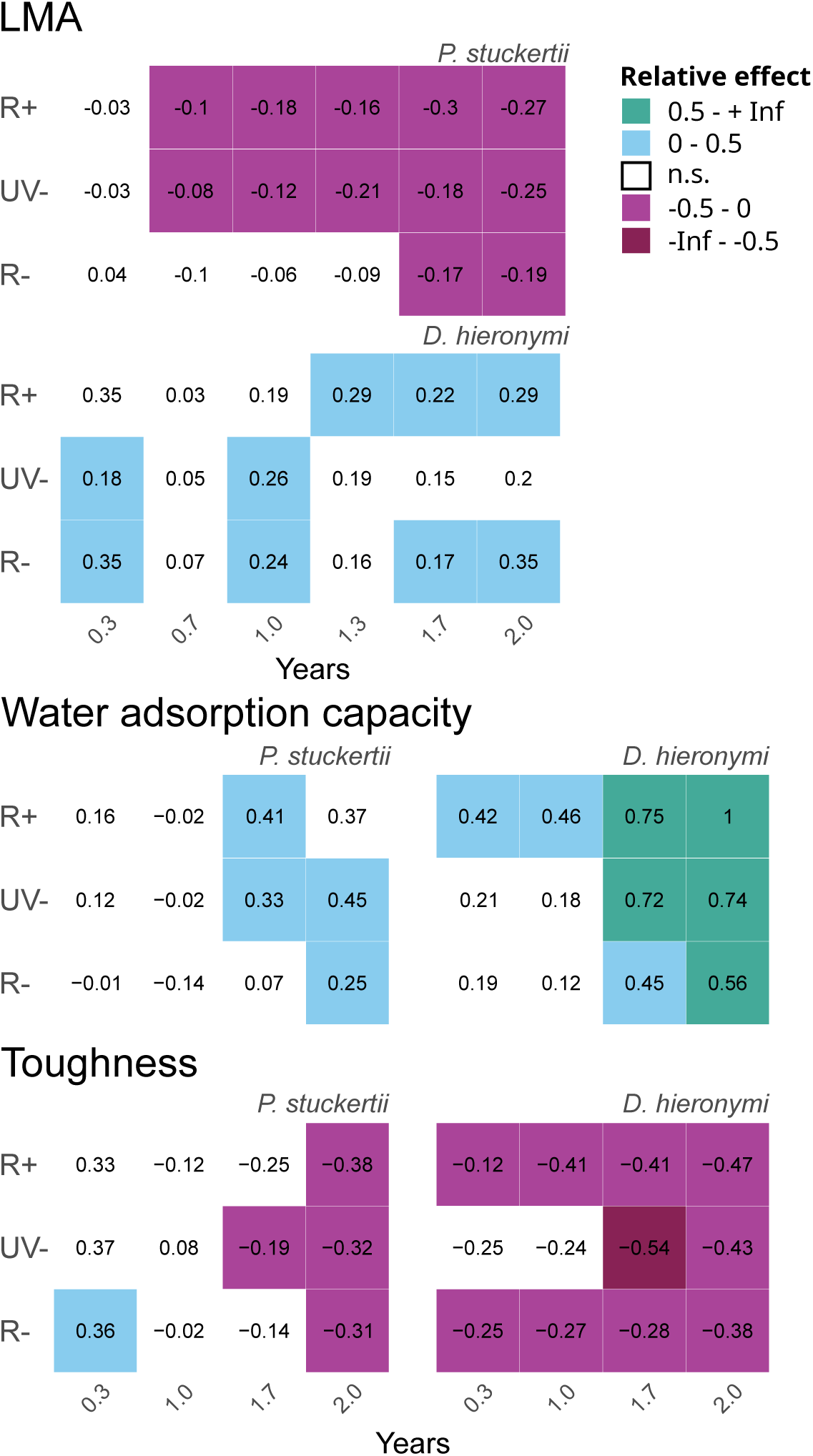
Relative effect of field decomposition on physical traits compared to initial values for both species under each filter. Colored squares denote significant differences for a t-test (*p <* 0.05). Colors represent direction of change and intensity: blue shades represent increases in trait values and pink shades represent decreases, while lighter shades represent low-moderate changes (0 – ±0.5) and darker shades represent bigger changes (±0.5 – infinite). LMA: leaf mass per area.

In contrast, *D. hieronymi* did not show a clear response in LMA to sunlight exposure. In terms of water adsorption capacity, this species responded to UV light at the beginning of the experiment with a 42% increase at 0.3 years (*p*: 0.005), and this pattern continued until 1 year (+46%; *p*: 0.02). All samples became more hydrophilic at the end of the experiment. Leaf toughness showed no clear response to sunlight exposure, although leaf toughness for this species also decreased over time. Finally, *D. hieronymi* had continuously higher leaching throughout the experiment compared to initial conditions but with no clear response to sunlight exposure (Appendix S1: Figure S3).

## Discussion

In this study, we set out to understand how important photodegradation was for C turnover of plant litter in a monsoonal mountain grassland. We first hypothesized that sunlight accelerates C cycling at this site and that this dynamic is modulated by seasonality. Our results confirm this hypothesis, although we found diverging patterns between the two species of dominant grasses. Moreover, we hypothesized that photofacilitation would occur during humid seasons due to sunlight exposure in the previous dry season. This was partially confirmed by results in *D. hieronymi*, but not in *P. stuckertii*, reinforcing how differently both species react to sunlight exposure. Our last hypothesis proposed that photodegradation produces changes in litter physical traits over time. We found partial support for this hypothesis, most clearly with changes in *P. stuckertii* ’s LMA and *D. hieronymi* ’s water adsorption capacity.

Generally, highly productive grasslands have been almost absent from photodegradation research (Austin and Ballaŕe, 2024). Overall, we demonstrate that photodegradation can considerably enhance plant litter decomposition rates in productive mountain grasslands. However, we did not expect to find such contrasting patterns for spectral dependence of photodegradation and response to sunlight exposure between the two species under study, given that most aspects of initial litter quality were similar (Figure 5). Most notably, *D. hieronymi* showed a 46% increase in mass loss after 1.3 years of UV-light exposure, while *P. stuckertii* had only a 15% increase in mass loss due to visible-light exposure at the same date (Figure 2). Other field experiments in grasslands with similar precipitation to our study site found either high (Butler et al., 2023) or intermediate (Brandt et al., 2010) effects of photodegradation. Butler et al. (2023) found an increase of up to 50% in mass loss with full sunlight exposure. Meanwhile, Brandt et al. (2010) found 17% increase in decomposition rates in one species but not in the other one. This could have been explained by the higher N concentration of the former species due to a confounded effect of biotic decomposition or even photofacilitation. We did not measure N concentration of our samples, but *D. hieronymi* has a slightly higher N concentration than *P. stuckertii* which might explain the bigger mass losses (Poca et al., 2014). These studies are not directly comparable, however, due to the fact that they did not include all the attenuation treatments (Brandt et al., 2010), or could not distinguish UV and visible light effects (Butler et al., 2023). Moreover, our study is the only one to assess photodegradation of standing dead litter in this type of grassland. Taken together, this makes comparisons difficult and highlights the need to include both sunlight spectra in future experiments. Photofacilitation can be defined as the increase in microbially mediated decomposition as a consequence of litter alterations due to the effect of photodegradation (Austin and Ballaŕe, 2024). We were able to infer accelerated biotic decomposition at our grassland site by measuring extracellular enzymatic activities, and again we found striking differences between the two species. The accumulated activity of the hydrolytic enzyme *β*-glucosidase did not significantly respond to light in *P. stuckertii*, but it did increase 50% under UV-light exposure in *D. hieronymi* after 2 years (Figure 2). This supports the idea that identity of dominant species is important in determining both magnitude and direction in biogeochemical processes (Fan et al., 2024). Connecting back to studies in similar grasslands, Brandt et al. (2010) did not find any photodegradation effect on *β*-glucosidase over the course of their study. Both experiments lasted for about 2 years, but there were seasonality differences between the monsoonal regime (this study) and the Mediterranean climate in Brandt et al. (2010). We were able to measure photofacilitation towards the end of the experiment, probably because this coincided with the end of the humid season. Meanwhile, their study ended during the dry season, when biotic decomposition is typically reduced. In this regard, a study with a similar design to ours in a Mediterranean open woodland did find photofacilitation effects of *β*-glucosidase activity (Berenstecher et al., 2020). Their study ended after the wet season, just like ours, which possibly allowed for the effects of photodegradation to accumulate over time, finally boosting biotic decomposition once humidity became available. This suggests that seasonal variation of rainfall and temperature can determine how abiotic and biotic decomposition mechanisms unfold.

It is clear that precipitation seasonality plays a big role in determining the directions and magnitude of photodegradative effects in grassland ecosystems. Aside from the evidence of photofacilitation mentioned above, we were able to detect other effects of seasonality on decomposition at this grassland. This was possible due to our explicit experiment design that was customized to closely follow wet and dry seasons, following Berenstecher et al. (2020). First, we detected a halt in decomposition during the second dry season (between 1 and 1.3 years), except for *D. hieronymi* under full sun treatment (Figure 3). This suggests that biotic activity was negligible during this period of low humidity, but that UV radiation exposure maintained C losses for that species. Moreover, this direct photodegradation effect probably caused the apparent photofacilitation found at the subsequent humid season, as previously discussed. Next, we found that the order of wet and dry seasons affected decomposition rates of *P. stuckertii* but not of *D. hieronymi* (Figure 4). Cumulatively, our results suggest that the former responds more strongly to rainfall seasonality, probably due to its higher water adsorption capacity (Figure 5), while the latter is more responsive to solar radiation.

Only two photodegradation studies used a similar temporal design to evaluate seasonality effects (Berenstecher et al., 2020; Li et al., 2024). Berenstecher et al. (2020) found that samples under full sunlight after a year decomposed faster when they started in the dry season compared to starting in the humid season, similar to *P. stuckertii* in this study (Figure 4). Li et al. (2024), additionally, studied photodegradation of forest litter with seasonal snow cover. They found that litter that was exposed to sunlight at the start during the snow-free autumn decomposed faster than samples that started during the snow-covered winter. Evidently, although with differences between ecosystem types, the alternation of precipitation and temperature seasons modulates the effects of photodegradation, and more studies should apply such experimental design to expand our understanding of this interaction.

Decomposition not only entails organic mass loss, but also chemical and physical changes as a response to biotic and abiotic drivers (Prescott and Vesterdal, 2021). It is important to study these changes through time because they in turn affect the trajectory of the decomposition process and determine the chemical quality of the remaining organic matter entering the soil (Cotrufo et al., 2015). However, most studies focus on changes in chemical traits over time (Brandt et al., 2010; Uselman et al., 2011; Méndez et al., 2019), and nearly no photodegradation studies have focused on physical traits. We detected a clear decrease in LMA as an effect of blue and green light on *P. stuckertii* (Figure 6). Only one previous study reported a similar decrease in LMA with exposure to UV light after 5 months (Gaxiola and Armesto, 2015). A likely explanation for this result is that leaf mass was lost with little changes in leaf area, resulting in thinner litter and thus decreased LMA. We also detected a clear increase in water adsorption capacity in both species (Figure 6), a proxy of litter hydrophilia (Talhelm and Smith, 2018). This response to solar radiation could be a consequence of the degradation of hydrophobic leaf cuticles (Logan et al., 2022), and other hydrophobic structural leaf compounds like lignin that respond strongly to photodegradation (Austin and Ballaŕe, 2010; Wang et al., 2024). Interestingly, the changes in these two litter traits might be connected to lignin losses through photodegradation, which would explain why litter became lighter and more hydrophilic over time (Austin and Ballaŕe, 2024). Considering physical litter traits affect decomposition dynamics and in turn those traits change over time with decomposition, it becomes evident that the relationship between them adjusts dynamically over time and determines the fate of litter C (Sun et al., 2022).

## Conclusions

Our study shows clear evidence that the effect of photodegradation is not limited to arid environments, but it also affects mesic and highly productive ecosystems like grasslands. We show proof that in these ecosystems a considerable amount of mass loss happens in the air before litter even touches the ground, both through direct abiotic and biotic contributions of solar radiation. Grasslands are estimated to cover 22.8% of the global land surface, representing a substantial proportion of terrestrial area (MacDougall et al., 2026). In these grass-dominated ecosystems, standing dead biomass constitutes a large portion of total biomass (Sarmiento, 1992; Yang et al., 2024). Hence, we can project that wherever grasslands with marked seasonality are found and standing dead biomass persists for long periods, photodegradation will be an important driver of C cycling (Keiser and Nieland, 2025; Yang et al., 2025). Further, focusing on grasslands with precipitations ruled by a monsoon, the impact of photodegradation could potentially be even larger. Recently, research has shown that in global drylands seasonality is an important driver of decomposition, with faster decomposition in sites under monsoonal compared to Mediterranean climates (Siebenhart et al., 2025). The coincidence of peaks in temperature and precipitation can boost microbial processes, including interactive effects with abiotic mechanisms like photofacilitation. Photodegradation has many implications in the terrestrial C cycle (Austin and Ballaŕe, 2024), from direct C emissions through photomineralization to changes in litter quality that affect the afterlife of plant-derived organic matter in soils. Taking all into consideration, it becomes clear that the study of photodegradation in productive grasslands with monsoonal climates should be a priority for a better understanding of the terrestrial C cycle.

## Acknowledgements

We would like to thank Quebrada del Condorito National Park, especially Fernanda Fabbio, for facilitating our field experiment; Cecilia Palmieri for granting us access to her property during field trips; Jośe Luis Lois, Franco Fernández and Lucio Biancari for their assistance during field work; Laura Ventura for her assistance in the laboratory; Gonzalo Arias and Lucas Enrico from IMBiV for their help with leaf toughness measurements.

## Author Contributions

Agustín Sarquis: Conceptualization, Methodology, Formal Analysis, Investigation, Writing - Original Draft, Visualization, Funding Acquisition, Project Administration. Ignacio A. Siebenhart: Investigation, Writing - Review and Editing. Marcela S. Méndez: Investigation, Writing - Review and Editing. Amy T. Austin: Conceptualization, Methodology, Funding Acquisition, Resources, Supervision, Writing - Review and Editing.

## Conflict of Interest Statement

We declare that none of the authors have any conflict of interest.

## Notes

### Competing Interest Statement

The authors have declared no competing interest.

### Funding

- University of Buenos Aires
- Fondo para la Investigación Científica y Tecnológica (FONCYT; Agencia I+D+I, Argentina)
- New Phytologist Foundation
- Fondation L’Oréal for Women in Science, France
- Alexander Von Humboldt Stiftung, Germany
- Neotropical Grassland Conservancy, US

## References

Administración de Parques Nacionales (2021). Disposicíon aprobatoria de las Cargas Ganaderas del Programa de Herbivoŕıa Doméstica del Parque Nacional Quebrada del Condorito: DI-2021-44637752-APN-DRC#APNAC. Technical report.

Almagro, M., F. T. Maestre, J. Martínez-Ĺopez, E. Valencia, and A. Rey (2015). Climate change may reduce litter decomposition while enhancing the contribution of photodegradation in dry perennial Mediterranean grasslands. Soil Biol. Biochem. 90, 214–223.

Almagro, M., J. Martínez-Ĺopez, F. T. Maestre, and A. Rey (2017). The Contribution of Photodegradation to Litter Decomposition in Semiarid Mediterranean Grasslands Depends on its Interaction with Local Humidity Conditions, Litter Quality and Position. Ecosystems 20 (3), 527–542.

Araujo, P. I., A. A. Grasso, A. González-Arzac, M. S. Méndez, and A. T. Austin (2022, jun). Sunlight and soil biota accelerate decomposition of crop residues in the Argentine Pampas. Agric. Ecosyst. Environ. 330, 107908.

Austin, A. T. and C. L. Ballaŕe (2010). Dual role of lignin in plant litter decomposition in terrestrial ecosystems. Proc. Natl. Acad. Sci. 107 (10), 4618–4622.

Austin, A. T. and C. L. Ballaŕe (2024, nov). Photodegradation in terrestrial ecosystems. New Phytol. 244 (3), 769–785.

Austin, A. T., M. S. Méndez, and C. L. Ballaŕe (2016). Photodegradation alleviates the lignin bottleneck for carbon turnover in terrestrial ecosystems. Proc. Natl. Acad. Sci. 113 (16), 4392–4397.

Austin, A. T., O. E. Sala, and R. B. Jackson (2006, dec). Inhibition of Nitrification Alters Carbon Turnover in the Patagonian Steppe. Ecosystems 9 (8), 1257–1265.

Austin, A. T. and L. Vivanco (2006). Plant litter decomposition in a semi-arid ecosystem controlled by photodegradation. Nature 442 (7102), 555–558.

Baker, N. R. and S. D. Allison (2015, jul). Ultraviolet photodegradation facilitates microbial litter decomposition in a Mediterranean climate. Ecology 96 (7), 1994–2003.

Ball, B. A., M. P. Christman, and S. J. Hall (2019, jan). Nutrient dynamics during photodegradation of plant litter in the Sonoran Desert. J. Arid Environ. 160 (January 2018), 1–10.

Berenstecher, P., L. Vivanco, and A. T. Austin (2022). Summer sunlight impacts carbon turnover in a spatially heterogeneous Patagonian woodland. Plant Soil (0123456789).

Berenstecher, P., L. Vivanco, L. I. Pérez, C. L. Ballaŕe, and A. T. Austin (2020, jul). Sunlight Doubles Aboveground Carbon Loss in a Seasonally Dry Woodland in Patagonia. Curr. Biol. 30, 1–9.

Bradford, M. A., G. F. Veen, A. Bonis, E. M. Bradford, A. T. Classen, J. H. C. Cornelissen, T. W. Crowther, J. R. De Long, G. T. Freschet, P. Kardol, M. Manrubia-Freixa, D. S. Maynard, G. S. Newman, R. S. P. Logtestijn, M. Viketoft, D. A. Wardle, W. R. Wieder, S. A. Wood, and W. H. van der Putten (2017, nov). A test of the hierarchical model of litter decomposition. Nat. Ecol. Evol. 1 (12), 1836–1845.

Brandt, L. A., C. Bohnet, and J. Y. King (2009, jun). Photochemically induced carbon dioxide production as a mechanism for carbon loss from plant litter in arid ecosystems. J. Geophys. Res. Biogeosciences 114 (G2), 1–13.

Brandt, L. A., J. Y. King, S. E. Hobbie, D. G. Milchunas, and R. L. Sinsabaugh (2010, aug). The Role of Photodegradation in Surface Litter Decomposition Across a Grassland Ecosystem Precipitation Gradient. Ecosystems 13 (5), 765–781.

Brandt, L. A., J. Y. King, and D. G. Milchunas (2007, oct). Effects of ultraviolet radiation on litter decomposition depend on precipitation and litter chemistry in a shortgrass steppe ecosystem. Glob. Chang. Biol. 13 (10), 2193–2205.

Breuil, C. and J. Saddler (1985, jul). Comparison of the 3,5-dinitrosalicylic acid and Nelson-Somogyi methods of assaying for reducing sugars and determining cellulase activity. Enzyme Microb. Technol. 7 (7), 327–332.

Butler, F. E. B., M. K. Good, J. W. Morgan, and N. L. Schultz (2023, dec). Relative contribution of photodegradation to litter breakdown in Australian grasslands. Ecol. Evol. 13 (12).

Cabido, M., R. Breimer, and G. Vega (1987). Plant Communities and Associated Soil Types in a High Plateau of the Cordoba Mountains, Central Argentina. Mt. Res. Dev. 7, 25–42.

Cabrera, F., P. I. Araujo, and L. Vivanco (2026, may). Photodegradation and microbial decomposition of soybean and maize crop residues before and after harvest. Agric. Ecosyst. Environ. 401, 110293.

Cadisch, G. and K. Giller (1997). Driven by nature: plant litter quality and decomposition. Wallingford, UK: CABI Publishing.

Chen, F. and R. A. Dixon (2007, jul). Lignin modification improves fermentable sugar yields for biofuel production. Nat. Biotechnol. 25 (7), 759–761.

Cingolani, A. M., M. Poca, M. A. Giorgis, M. V. Vaieretti, D. E. Gurvich, J. I. Whitworth-Hulse, and D. Renison (2015). Water provisioning services in a seasonally dry subtropical mountain: Identifying priority landscapes for conservation. J. Hydrol. 525, 178–187.

Cingolani, A. M., M. V. Vaieretti, M. A. Giorgis, N. La Torre, J. I. Whitworth-Hulse, and D. Renison (2013). Can livestock and fires convert the sub-tropical mountain rangelands of central Argentina into a rocky desert? Rangel. J. 35 (3), 285–297.

Cingolani, A. M., M. V. Vaieretti, M. A. Giorgis, M. Poca, P. A. Tecco, and D. E. Gurvich (2014). Can livestock grazing maintain landscape diversity and stability in an ecosystem that evolved with wild herbivores? Perspect. Plant Ecol. Evol. Syst. 16 (4), 143–153.

Cornwell, W. K., J. H. C. Cornelissen, K. Amatangelo, E. Dorrepaal, V. T. Eviner, O. Godoy, S. E. Hobbie, B. Hoorens, H. Kurokawa, N. Pérez-Harguindeguy, H. M. Quested, L. S. Santiago, D. A. Wardle, I. J. Wright, R. Aerts, S. D. Allison, P. van Bodegom, V. Brovkin, A. Chatain, T. V. Callaghan, S. Díaz, E. Garnier, D. E. Gurvich, E. Kazakou, J. A. Klein, J. Read, P. B. Reich, N. A. Soudzilovskaia, M. V. Vaieretti, and M. Westoby (2008, oct). Plant species traits are the predominant control on litter decomposition rates within biomes worldwide. Ecol. Lett. 11 (10), 1065–1071.

Cotrufo, M. F. and J. M. Lavallee (2022). Soil organic matter formation, persistence, and functioning: A synthesis of current understanding to inform its conservation and regeneration. In Adv. Agron, Volume 172, pp. 1–66. Elsevier Inc.

Cotrufo, M. F., J. L. Soong, A. J. Horton, E. E. Campbell, M. L. Haddix, D. H. Wall, and W. J. Parton (2015). Formation of soil organic matter via biochemical and physical pathways of litter mass loss. Nat. Geosci. 8 (10), 776–779.

Day, T. A. and M. S. Bliss (2019, dec). A spectral weighting function for abiotic photodegradation based on photochemical emission of CO2 from leaf litter in sunlight. Biogeochemistry 146 (2), 173–190.

Day, T. A. and M. S. Bliss (2020, nov). Solar Photochemical Emission of CO2 From Leaf Litter: Sources and Significance to C Loss. Ecosystems 23 (7), 1344–1361.

Day, T. A., M. S. Bliss, A. R. Tomes, C. T. Ruhland, and R. Gúenon (2018). Desert leaf litter decay: Coupling of microbial respiration, water-soluble fractions and photodegradation. Glob. Chang. Biol. 24 (11), 5454–5470.

Day, T. A., R. Gúenon, and C. T. Ruhland (2015). Photodegradation of plant litter in the Sonoran Desert varies by litter type and age. Soil Biol. Biochem. 89, 109–122.

Day, T. A., J. M. Urbine, and M. S. Bliss (2022, feb). Supplemental precipitation accelerates decay but only in photodegraded litter and implications that sunlight promotes leaching loss. Biogeochemistry 158 (1), 113–129.

Day, T. A., E. T. Zhang, and C. T. Ruhland (2007, oct). Exposure to solar UV-B radiation accelerates mass and lignin loss of Larrea tridentata litter in the Sonoran Desert. Plant Ecol. 193 (2), 185–194.

DuBois, M., K. A. Gilles, J. K. Hamilton, P. A. Rebers, and F. Smith (1956, mar). Colorimetric Method for Determination of Sugars and Related Substances. Anal. Chem. 28 (3), 350–356.

Erdenebileg, E., X. Ye, C. Wang, Z. Huang, G. Liu, and J. H. Cornelissen (2018, sep). Positive and negative effects of UV irradiance explain interaction of litter position and UV exposure on litter decomposition and nutrient dynamics in a semi-arid dune ecosystem. Soil Biol. Biochem. 124 (January), 245–254.

Fan, B., Z. Gong, X. Xin, Y. Liu, L. He, Y. Gao, A. Ren, and N. Zhao (2024, feb). Both evenness and dominant species identity have effects on litter decomposition. Ecol. Evol. 14 (2), 1–13.

Foereid, B., J. Bellarby, W. Meier-Augenstein, and H. Kemp (2010). Does light exposure make plant litter more degradable? Plant Soil.

Gallo, M. E., A. Porras-Alfaro, K. J. Odenbach, and R. L. Sinsabaugh (2009). Photoacceleration of plant litter decomposition in an arid environment. Soil Biol. Biochem. 41 (7), 1433–1441.

Gallo, M. E., R. L. Sinsabaugh, and S. E. Cabaniss (2006). The role of ultraviolet radiation in litter decomposition in arid ecosystems. Appl. Soil Ecol. 34 (1), 82–91.

Gaxiola, A. and J. J. Armesto (2015). Understanding litter decomposition in semiarid ecosystems: linking leaf traits, UV exposure and rainfall variability. Front. Plant Sci. 6.

Gholz, H. L., D. A. Wedin, S. M. Smitherman, M. E. Harmon, and W. J. Parton (2000, oct). Long-term dynamics of pine and hardwood litter in contrasting environments: toward a global model of decomposition. Glob. Chang. Biol. 6 (7), 751–765.

Ghose, T. K. (1987, jan). Measurement of cellulase activities. Pure Appl. Chem. 59 (2), 257–268.

Gibson, D. J. (2008, oct). Grasses and Grassland Ecology. Oxford University PressOxford.

Harmon, M., K. Nadelhoffer, and J. Blair (1999). Measuring decomposition, nutrient turnover, and stores in plant litter. In G. Robertson, D. Coleman, C. Bledsoe, and P. Sollins (Eds.), Stand. Soil Methods Long-Term Ecol. Res., pp. 202–240. Oxford: Oxford University Press.

Henry, H. A., K. Brizgys, and C. B. Field (2008). Litter decomposition in a California annual grassland: Interactions between photodegradation and litter layer thickness. Ecosystems 11 (4), 545–554.

Hosseiniaghdam, E., H. Yang, M. Mamo, M. Kaiser, W. H. Schacht, K. M. Eskridge, and G. O. Abagandura (2023, feb). Effects of litter placement, soil moisture and temperature on soil carbon dioxide emissions in a sandy grassland soil. Grassl. Sci. (November 2021), 1–10.

Huang, G., H. Zhao, and Y. Li (2017). Litter decomposition in hyper-arid deserts: Photodegradation is still important. Sci. Total Environ. 601-602, 784–792.

Keiser, A. D. and M. A. Nieland (2025, dec). Interactions Are Key to Accurately Estimating the Impact of Photodegradation Across Grassland Ecosystems. Glob. Chang. Biol. 31 (12).

Keiser, A. D., R. Warren, T. Filley, and M. A. Bradford (2021, apr). Signatures of an abiotic decomposition pathway in temperate forest leaf litter. Biogeochemistry 153 (2), 177–190.

King, J. Y., L. A. Brandt, and E. C. Adair (2012). Shedding light on plant litter decomposition: Advances, implications and new directions in understanding the role of photodegradation. Biogeochemistry 111 (1-3), 57–81.

Kommedal, E. G., C. F. Angeltveit, L. J. Klau, I. Ayuso-Ferńandez, B. Arstad, S. G. Antonsen, Y. Stenstrøm, D. Ekeberg, F. Gírio, F. Carvalheiro, S. J. Horn, F. L. Aachmann, and V. G. H. Eijsink (2023, feb). Visible light-exposed lignin facilitates cellulose solubilization by lytic polysaccharide monooxygenases. Nat. Commun. 14 (1), 1063.

Lee, H., T. Rahn, and H. Throop (2012, mar). An accounting of C-based trace gas release during abiotic plant litter degradation. Glob. Chang. Biol. 18 (3), 1185–1195.

Lejoly, J. D. M., K. Mason-Jones, and G. F. C. Veen (2026, feb). A soil food web approach to integrate soil fauna into multitrophic biogeochemistry. Commun. Earth Environ..

Li, X., Y. Wang, J. Zhang, T. M. Robson, H. Kurokawa, H. Peng, L. Zhou, D. Yu, J. Deng, and Q.-W. Wang (2024, jun). Autumn sunlight promotes aboveground carbon loss in a temperate mixed forest. Ecol. Process. 13 (1), 48.

Logan, J. R., P. Barnes, and S. E. Evans (2022, jul). Photodegradation of plant litter cuticles enhances microbial decomposition by increasing uptake of non-rainfall moisture. Funct. Ecol. 36 (7), 1727–1738.

Ma, Z., W. Yang, F. Wu, and B. Tan (2017, apr). Effects of light intensity on litter decomposition in a subtropical region. Ecosphere 8 (4).

MacDougall, A. S., B. Vanzant, J. Sulik, S. Bagchi, D. Naidu, T. O. Muraina, E. W. Seabloom, E. T. Borer, P. Wilfahrt, I. Slette, J. L. Hierro, D. E. Pearson, M. Abedi, M. Akasaka, J. Alberti, A. Aleksanyan, A. A. Amisu, T. M. Anderson, C. A. Arnillas, M. Ayer, J. D. Bakker, S. Basant, S. Basto, L. Biederman, K. J. Bloodworth, F. Boscutti, E. H. Boughton, C. M. Bruschetti, H. L. Buckley, Y. M. Buckley, M. N. Bugalho, M. C. Caldeira, G. Campetella, N. Cannone, M. Carbognani, C. Carbutt, M. A. Carniello, M. Cervellini, T. Chaudhary, Q. Chen, A. T. Clark, S. Cousins, M. Dalle Fratte, N. J. Day, B. Déak, J. Dietrich, A. Dixon, N. Eisenhauer, K. J. Elgersma, O. Eren, A. Eskelinen, C. Estrada, P. A. Fay, G. Fayvush, K. C. Flynn, D. Garćıa Meza, D. Gargano, L. Gherardi, N. T. Girkin, L. Gonźalez, P. Graff, L. W. C. Hagenberg, A. H. Halbritter, N. A. Havrilchak, N. Herdoiza, E. Hersch-Green, K. Hopping, A. Jentsch, S. O. Jimoh, J. Kerby, K. Kirkman, J. M. H. Knops, S. E. Koerner, A. Koltz, K. J. Komatsu, B. I. Koura, S. Kruse, L. Laanisto, L. S. Lannes, W. Li, M. Liang, A. Lkhagva, L. López-Olmedo, P. Lorenzo, C. J. Lortie, A. Loydi, W. Luo, P. Macek, F. Malfasi, P. Mariotte, J. P. Martina, A. Martínez-Blancas, H. Martinson, C. Martorell, J. A. Meave, S. Medina-Villar, K. Z. Mganga, J. Monsimet, A. N. Nerlekar, S. Niu, T. Ohlert, I. Oliveras Menor, G. R. Oñatibia, Y. K. Ortega, B. Osborne, S. Palpurina, J. Pascual, S. C. Pennings, E. Pérez-Garćıa, P. L. Peri, M. Petit Bon, A. Petraglia, F. Pijcke, S. M. Prober, R. E. Quiroga, J. I. Ramirez, S. Reed, B. H. P. Rosado, C. Roscher, D. W. Rowley, I. Sereda, D. M. Small, N. G. Smith, Y. Song, C. Stevens, L. E. Suarez Jimenez, M. te Beest, M. Tedder, R. S. Terry, K. S. Thornton, D. Tian, G. Titcomb, O. Valkó, G. F. ‘Ciska’ Veen, R. Virtanen, E. A. R. Welti, G. R. Wheeler, A. A. Wolf, P. Wolff, A. L. Young, H. S. Young, L. H. Zeglin, K. Zhu, S. Zong, and M. B. Siewert (2026, jan). The global extent of the grassland biome and implications for the terrestrial carbon sink. Nat. Ecol. Evol. 10 (2), 246–257.

Marinho, O. A., L. A. Martinelli, P. J. Duarte-Neto, E. A. Mazzi, and J. Y. King (2020, may). Photodegradation influences litter decomposition rate in a humid tropical ecosystem, Brazil. Sci. Total Environ. 715, 136601.

Masubelele, M. L. and W. Bond (2022, sep). Grass species litter have varied trait response to the photodegradation and microbial decomposition in tropical savanna grasslands, South Africa. Ann. Environ. Sci. Toxicol. 6 (1), 054–062.

Mazza, C. A., H. E. Boccalandro, C. V. Giordano, D. Battista, A. L. Scopel, and C. L. Ballaré (2000, jan). Functional Significance and Induction by Solar Radiation of Ultraviolet-Absorbing Sunscreens in Field-Grown Soybean Crops. Plant Physiol. 122 (1), 117–126.

Méndez, M. S., C. L. Ballaŕe, and A. T. Austin (2022, sep). Dose–responses for solar radiation exposure reveal high sensitivity of microbial decomposition to changes in plant litter quality that occur during photodegradation. New Phytol. 235 (5), 2022–2033.

Méndez, M. S., M. L. Martinez, P. I. Araujo, and A. T. Austin (2019, dec). Solar radiation exposure accelerates decomposition and biotic activity in surface litter but not soil in a semiarid woodland ecosystem in Patagonia, Argentina. Plant Soil 445 (1-2), 483–496.

Moorhead, D. L. and T. Callaghan (1994). Effects of increasing ultraviolet B radiation on decomposition and soil organic matter dynamics: a synthesis and modelling study. Biol. Fertil. Soils.

Mudrák, O., S^̌^. Angst, G. Angst, H. Veseĺa, R. Schnablová, T. Herben, and J. Frouz (2023, aug). Ecological significance of standing dead phytomass: Marcescence as a puzzle piece to the nutrient cycle in temperate ecosystems. J. Ecol. (May 2022), 1–12.

Olson, J. S. (1963, apr). Energy Storage and the Balance of Producers and Decomposers in Ecological Systems. Ecology 44 (2), 322–331.

Pancotto, V. A., O. E. Sala, M. Cabello, N. I. Lopez, T. Matthew Robson, C. L. Ballare, M. M. Caldwell, and A. L. Scopel (2003, oct). Solar UV-B decreases decomposition in herbaceous plant litter in Tierra del Fuego, Argentina: potential role of an altered decomposer community. Glob. Chang. Biol. 9 (10), 1465–1474.

Pérez-Harguindeguy, N., S. Díaz, E. Garnier, S. Lavorel, H. Poorter, P. Jaureguiberry, M. S. Bret-Harte, W. K. Cornwell, J. M. Craine, D. E. Gurvich, C. Urcelay, E. J. Veneklaas, P. B. Reich, L. Poorter, I. J. Wright, P. Ray, L. Enrico, J. G. Pausas, A. C. de Vos, N. Buchmann, G. Funes, F. Qúetier, J. G. Hodgson, K. Thompson, H. D. Morgan, H. ter Steege, L. Sack, B. Blonder, P. Poschlod, M. V. Vaieretti, G. Conti, A. C. Staver, S. Aquino, and J. H. C. Cornelissen (2013, may). New handbook for standardised measurement of plant functional traits worldwide. Aust. J. Bot. 61 (3), 167.

Poca, M., A. M. Cingolani, D. E. Gurvich, V. Saur Palmieri, and G. Bertone (2018). Water storage dynamics across different types of vegetated patches in rocky highlands of central Argentina. Ecohydrology 11 (7), 1–14.

Poca, M., N. Pérez Harguindeguy, M. V. Vaieretti, and A. M. Cingolani (2014, aug). Descomposición y calidad físico-qúımica foliar de 24 especies dominantes de los pastizales de altura de las sierras de Córdoba, Argentina. Ecol. Austral 24 (2), 249–257.

Prescott, C. E. and L. Vesterdal (2021, oct). Decomposition and transformations along the continuum from litter to soil organic matter in forest soils. For. Ecol. Manage. 498 (July), 119522.

Pucheta, E., M. Cabido, S. Díaz, and G. Funes (1998). Floristic composition, biomass, and aboveground net plant production in grazed and protected sites in a mountain grassland of central Argentina. Acta Oecologica 19 (2), 97–105.

R Core Team (2020). R: A language and environment for statistical computing.

Ruhland, C. T. and P. T. Fraley (2023, dec). The influence of initial phenolic content and UV-screening effectiveness on abiotic photodegradation of Wyoming big sagebrush litter collected along an elevation gradient. J. Arid Environ. 219 (October), 105084.

Rutledge, S., D. I. Campbell, D. Baldocchi, and L. A. Schipper (2010). Photodegradation leads to increased carbon dioxide losses from terrestrial organic matter. Glob. Chang. Biol. 16, 3065–3074.

Sarmiento, G. (1992). Adaptive strategies of perennial grasses in South American savannas. Journal of Vegetation Science 3 (3), 325–336.

Schade, G. W., R.-M. Hofmann, and P. J. Crutzen (2012). CO emissions from degrading plant matter. Tellus B Chem. Phys. Meteorol..

Schneider, C. A., W. S. Rasband, and K. W. Eliceiri (2012, jul). NIH Image to ImageJ: 25 years of image analysis. Nat. Methods 9 (7), 671–5.

Siebenhart, I. A., P. M. Tognetti, A. Sarquis, L. Biancari, C. L. Ballaŕe, and A. T. Austin (2025). Plant litter decomposition in global drylands is better predicted by precipitation seasonality and temperature than by aridity. bioRxiv.

Sinsabaugh, R. L., M. J. King, H. P. Collins, P. E. Yeager, and S. O. Petersen (1999). Characterizing Soil Microbial Communities. In P. D. Robertson, D. C. Coleman, C. S. Bledsoe, and P. Sollins (Eds.), Stand. Soil Methods Long-Term Ecol. Res. New York: Oxford University Press.

Sun, Z., P. Tian, X. Zhao, Y. Wang, S. Wang, X. Fang, Q. Wang, and S. Liu (2022). Temporal shifts in the explanatory power and relative importance of litter traits in regulating litter decomposition. For. Ecosyst. 9 (July), 100072.

Talhelm, A. F. and A. M. S. Smith (2018, apr). Litter moisture adsorption is tied to tissue structure, chemistry, and energy concentration. Ecosphere 9 (4), e02198.

Uselman, S. M., K. A. Snyder, R. R. Blank, and T. J. Jones (2011, jun). UVB exposure does not accelerate rates of litter decomposition in a semi-arid riparian ecosystem. Soil Biol. Biochem. 43 (6), 1254–1265.

Vaieretti, M. V., A. M. Cingolani, N. Pérez Harguindeguy, D. E. Gurvich, and M. Cabido (2010). Does decomposition of standard materials differ among grassland patches maintained by livestock? Austral Ecol. 35 (8), 935–943.

van Asperen, H., T. Warneke, S. Sabbatini, G. Nicolini, D. Papale, and J. Notholt (2015, jul). The role of photo- and thermal degradation for CO 2 and CO fluxes in an arid ecosystem. Biogeosciences 12 (13), 4161–4174.

Van Soest, P., J. Robertson, and B. Lewis (1991, oct). Methods for Dietary Fiber, Neutral Detergent Fiber, and Nonstarch Polysaccharides in Relation to Animal Nutrition. J. Dairy Sci. 74 (10), 3583–3597.

von Müller, A. R., D. Renison, and A. M. Cingolani (2017). Cattle landscape selectivity is influenced by ecological and management factors in a heterogeneous mountain rangeland. Rangel. J. 39, 1–14.

Wang, B., S. An, C. Liang, Y. Liu, and Y. Kuzyakov (2021, nov). Microbial necromass as the source of soil organic carbon in global ecosystems. Soil Biol. Biochem. 162 (September), 108422.

Wang, J., S. Yang, B. Zhang, W. Liu, M. Deng, S. Chen, and L. Liu (2017, oct). Temporal dynamics of ultraviolet radiation impacts on litter decomposition in a semi-arid ecosystem. Plant Soil 419 (1-2), 71–81.

Wang, P., Y. Liu, B. Zhang, L. Li, L. Lin, X. Li, and Q. Zeng (2024, mar). The effect of litter decomposition mostly depends on seasonal variation of ultraviolet radiation rather than species in a hyper-arid desert. Front. Environ. Sci. 12 (March), 1–13.

Wu, Q., X. Ni, X. Sun, Z. Chen, S. Hong, B. Berg, M. Zheng, J. Chen, J. Zhu, L. Ai, Y. Zhang, and F. Wu (2025, feb). Substrate and climate determine terrestrial litter decomposition. Proc. Natl. Acad. Sci. 122 (7).

Yang, S., Z. Jia, P. Chang, Y. Wu, J. Huang, J. Wang, M. Deng, J. Su, S. Hong, Y. He, J. Zhu, P. Zhang, Y. Wang, X. Guo, Z. Zhang, Y. Zhang, S. Hu, J. He, S. Piao, and L. Liu (2025, aug). Significant Impact of UV Exposure on Litter Decomposition Across Diverse Climate Zones. Glob. Chang. Biol. 31 (8).

Yang, S., J. Wang, J. Su, Z. Peng, L. Guo, Y. Wu, P. Chang, Y. Wang, J. Huang, and L. Liu (2024, jul). Divergent Roles of UV Exposure and Microclimatic Conditions in the Decomposition of Standing and Soil Surface Litter in a Semi-Arid Steppe. J. Geophys. Res. Biogeosciences 129 (7), 1–15.

Yao, B., X. Kong, K. Tian, X. Zeng, W. Lu, L. Pang, S. Sun, and X. Tian (2024, jul). Initial Litter Chemistry and UV Radiation Drive Chemical Divergence in Litter during Decomposition. Microorganisms 12 (8), 1535.

Yao, B., X. Zeng, L. Pang, X. Kong, K. Tian, Y. Ji, S. Sun, and X. Tian (2022, aug). The Photodegradation of Lignin Methoxyl C Promotes Fungal Decomposition of Lignin Aromatic C Measured with 13C-CPMAS NMR. J. Fungi 8 (9), 900.

Zanne, A. E., H. Flores-Moreno, J. R. Powell, W. K. Cornwell, J. W. Dalling, A. T. Austin, A. T. Classen, P. Eggleton, K.-i. Okada, C. L. Parr, E. C. Adair, S. Adu-Bredu, M. A. Alam, C. Alvarez-Garzón, D. Apgaua, R. Araǵon, M. Ardon, S. K. Arndt, L. A. Ashton, N. A. Barber, J. Beaucĥene, M. P. Berg, J. Beringer, M. M. Boer, J. A. Bonet, K. Bunney, T. J. Burkhardt, D. Carvalho, D. Castillo-Figueroa, L. A. Cernusak, A. W. Cheesman, T. M. Cirne-Silva, J. R. Cleverly, J. H. C. Cornelissen, T. J. Curran, A. M. D’Angioli, C. Dallstream, N. Eisenhauer, F. Evouna Ondo, A. Fajardo, R. D. Fernandez, A. Ferrer, M. A. L. Fontes, M. L. Galatowitsch, G. Gonźalez, F. Gottschall, P. R. Grace, E. Granda, H. M. Griffiths, M. Guerra Lara, M. Hasegawa, M. M. Hefting, N. Hinko-Najera, L. B. Hutley, J. Jones, A. Kahl, M. Karan, J. A. Keuskamp, T. Lardner, M. Liddell, C. Macfarlane, C. Macinnis-Ng, R. F. Mariano, M. S. Méndez, W. S. Meyer, A. S. Mori, A. S. Moura, M. Northwood, R. Ogaya, R. S. Oliveira, A. Orgiazzi, J. Pardo, G. Peguero, J. Penuelas, L. I. Perez, J. M. Posada, C. M. Prada, T. P̌ŕıv̌etivý, S. M. Prober, J. Prunier, G. W. Quansah, V. Resco de Dios, R. Richter, M. P. Robertson, L. F. Rocha, M. A. Rúa, C. Sarmiento, R. P. Silberstein, M. C. Silva, F. F. Siqueira, M. G. Stillwagon, J. Stol, M. K. Taylor, F. P. Teste, D. Y. P. Tng, D. Tucker, M. Türke, M. D. Ulyshen, O. J. Valverde-Barrantes, E. van den Berg, R. S. P. van Logtestijn, G. F. C. Veen, J. G. Vogel, T. J. Wardlaw, G. Wiehl, C. Wirth, M. J. Woods, and P.-C. Zalamea (2022, sep). Termite sensitivity to temperature affects global wood decay rates. Science (80-.). 377 (6613), 1440–1444.

Zeballos, S. R., J. J. Cantero, M. A. Giorgis, A. T. R. Acosta, C. O. Núñez, M. V. Palchetti, D. S. Argibay, and M. R. Cabido (2024, oct). Classification of montane grasslands in central Argentina. Appl. Veg. Sci. 27 (4).

Zhang, D., D. Hui, Y. Luo, and G. Zhou (2008, jun). Rates of litter decomposition in terrestrial ecosystems: global patterns and controlling factors. J. Plant Ecol. 1 (2), 85–93.

