## Appendix S1 for "Photodegradation accelerates standing dead litter decomposition in monsoonal mountain grasslands of South America"

Authors: Agustín Sarquis, Ignacio A. Siebenhart, Marcela S. Méndez, Amy T. Austin.

Manuscript title: Photodegradation accelerates standing dead litter decomposition in monsoonal mountain grasslands of South America

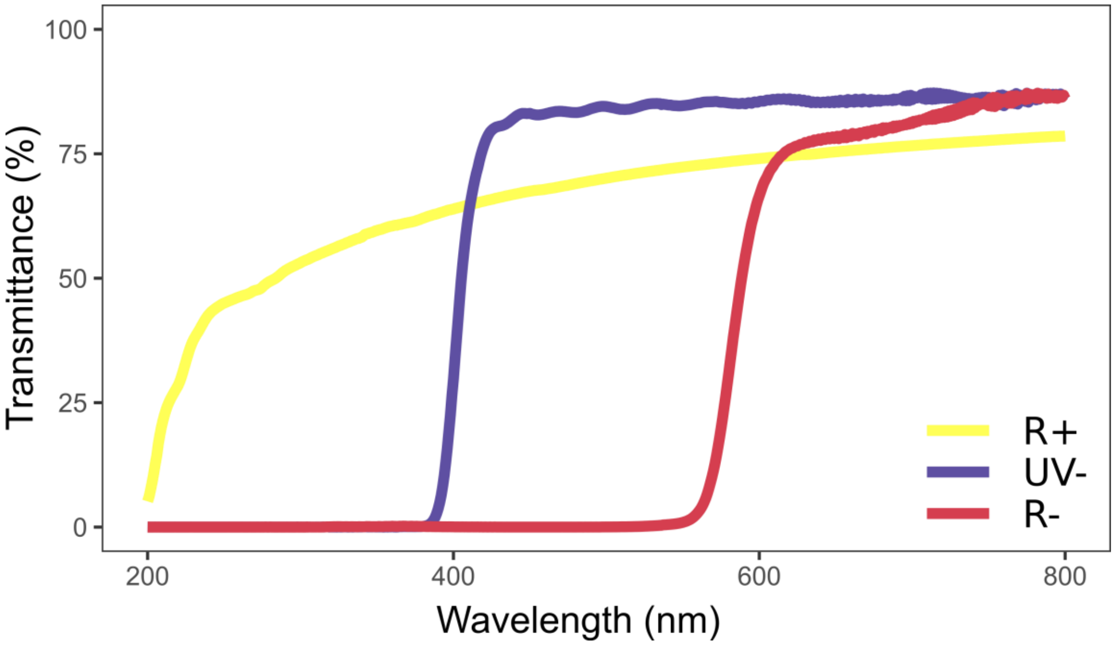


Figure S1: Filter transmittance (percentage of total incoming radiation, %). R+: 100 μm thickness polyethylene (280-800 nm; Ever Wear S. A.). UV-: 226 UV Costech filter (280-400 nm). R-: Rosco® Nº135 Deep Golden Amber filter (280-550 nm).

Table S1: Linear mixed models results for the temperature of envelopes in the field.

| Random effects | Variance | Standard Deviation | | | |
|---|---|---|---|---|---|
| Dates (intercept) | 30.45 | 5.52 | | | |
| Residuals | 5.85 | 2.42 | | | |
| Fixed Effects | Estimate | Standard Error | Degrees of freedom | t-value | p-value |
| R+ (Intercept) | 18.09 | 2.49 | 4.10 | 7.23 | 0.0017 ** |
| UV- | 17.42 | 0.44 | 173.01 | -1.53 | 0.13 |
| R- | 18.21 | 0.44 | 173.01 | 0.27 | 0.79 |
| Type III ANOVA | F | p | | | |
| Filter | 1.87 | 0.16 | | | |

Asterisks denote significant results: ** *p<0.01*.

R+: full radiation. UV-: UV blocked. R-: UV, blue and green light blocked.

Table S2: ANOVA results for remaining organic mass under the light attenuation treatment by species and year.

| Species / Group | Year | Degrees of freedom | $R^2$ | *p* |
|---|---|---|---|---|
| *P. stuckertii* | | | | |
| Winter Group 1 | 0.3 | 11 | 0.3 | 0.2 |
| | 1.0 | 11 | 0.0002 | 0.9 |
| | 1.3 | 11 | 0.6 | 0.006 ** |
| | 2.1 | 11 | 0.2 | 0.2 |
| Spring Group | 0.7 | 12 | 0.3 | 0.1 |
| | 1.0 | 12 | 0.1 | 0.5 |
| | 1.7 | 12 | 0.6 | 0.008 ** |
| Winter Group 2 | 0.3 | 11 | 0.2 | 0.3 |
| | 1.0 | 12 | 0.3 | 0.09 |
| *D. hieronymi* | | | | |
| Winter Group 1 | 0.3 | 8 | 0.4 | 0.1 |
| | 1.0 | 10 | 0.3 | 0.2 |
| | 1.3 | 10 | 0.7 | 0.003 ** |
| | 2.1 | 8 | 0.7 | 0.004 ** |
| Spring Group | 0.7 | 12 | 0.5 | 0.03 * |
| | 1.0 | 12 | 0.8 | 0.0001 *** |
| | 1.7 | 12 | 0.8 | < 0.0001 *** |
| Winter Group 2 | 0.3 | 11 | 0.1 | 0.5 |
| | 1.0 | 11 | 0.6 | 0.006 ** |

Asterisks denote significant results: *** $p<0.001$, ** $p<0.01$, * $p<0.05$.

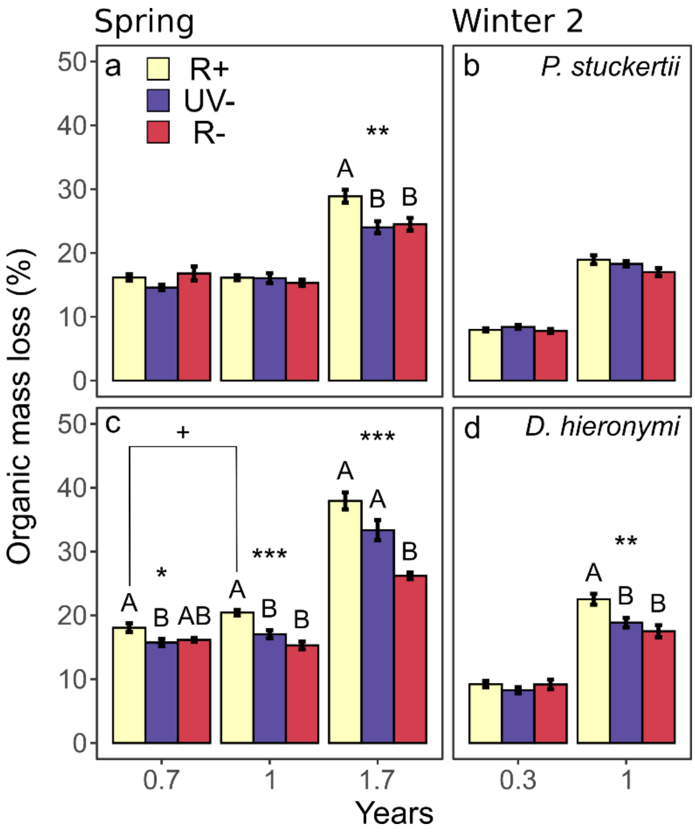


Figure S2: Organic mass (OM) loss (%) over time for both species of Spring and Winter Group 2. Asterisks denote significant results: *** *p<0.001*, ** *p<0.01*, * *p<0.05*. Upper-case letters denote differences between filters following Tukey HSD test. Bars represent mean values and standard errors. Results in this figure show OM mass loss, but statistical analyses were performed on remaining OM. R+: full radiation. UV-: UV blocked. R-: UV, blue and green light blocked.

Table S3: Accumulated β-glucosidase and phenol-oxidase enzymatic activities of litter over time (µ mol $g^{-1}$).

| Species | Group | Filter | Years | β-glucosidase | Phenol-oxidase |
|---|---|---|---|---|---|
| *P. stuckertii* | Winter 1 | R+ | 0.3 | 341 ± 59 | 0 ± 0 |
| | | | 1.0 | 3823 ± 965 | 38 ± 38 |
| | | | 1.3 | 15982 ± 2307 | 119 ± 95 |
| | | | 2.1 | 36425 ± 4955 | 239 ± 139 |
| | | UV- | 0.3 | 477 ± 33 | 0 ± 0 |
| | | | 1.0 | 5170 ± 772 | 36 ± 36 |
| | | | 1.3 | 20052 ± 3164 | 162 ± 94 |
| | | | 2.1 | 41022 ± 6129 | 119 ± 60 |
| | | R- | 0.3 | 488 ± 63 | 0 ± 0 |
| | | | 1.0 | 5243 ± 734 | 49 ± 49 |
| | | | 1.3 | 18542 ± 4234 | 160 ± 98 |
| | | | 2.1 | 29088 ± 2273 | 273 ± 112 |
| | Spring | R+ | 0.7 | 841 ± 130 | 0 ± 0 |
| | | | 1.0 | 8249 ± 889 | 11 ± 7 |
| | | | 1.7 | 23490 ± 2931 | 45 ± 27 |
| | | UV- | 0.7 | 1601 ± 378 | 17 ± 10 |
| | | | 1.0 | 8410 ± 1138 | 67 ± 20 |
| | | | 1.7 | 18727 ± 3310 | 121 ± 22 |
| | | R- | 0.7 | 1910 ± 220 | 13 ± 13 |
| | | | 1.0 | 10063 ± 770 | 51 ± 32 |
| | | | 1.7 | 22565 ± 2146 | 124 ± 28 |
| | Winter 2 | R+ | 0.3 | 961 ± 440 | 11 ± 4 |
| | | | 1.0 | 7703 ± 3364 | 53 ± 19 |
| | | UV- | 0.3 | 2089 ± 1016 | 10 ± 5 |
| | | | 1.0 | 11200 ± 4217 | 52 ± 24 |
| | | R- | 0.3 | 1632 ± 699 | 6 ± 3 |
| | | | 1.0 | 10217 ± 3379 | 32 ± 15 |
| *D. hieronyim* | Winter 1 | R+ | 0.3 | 589 ± 28 | 0 ± 0 |
| | | | 1.0 | 4779 ± 905 | 101 ± 47 |
| | | | 1.3 | 20808 ± 3533 | 300 ± 154 |
| | | | 2.1 | 54855 ± 3353 | 286 ± 89 |
| | | UV- | 0.3 | 645 ± 42 | 0 ± 0 |
| | | | 1.0 | 5234 ± 450 | 66 ± 46 |
| | | | 1.3 | 17468 ± 1760 | 270 ± 123 |
| | | | 2.1 | 37997 ± 5718 | 518 ± 195 |
| | | R- | 0.3 | 633 ± 67 | 0 ± 0 |
| | | | 1.0 | 6181 ± 590 | 67 ± 32 |

| | | | | |
|---|---|---|---|---|
| | | 1.3 | 18674 ± 1499 | 329 ± 103 |
| | | 2.1 | 36176 ± 4001 | 510 ± 170 |
| Spring | R+ | 0.7 | 1443 ± 357 | 0 ± 0 |
| | | 1.0 | 11887 ± 1094 | 43 ± 34 |
| | | 1.7 | 29866 ± 3417 | 181 ± 100 |
| | UV- | 0.7 | 1486 ± 362 | 5 ± 3 |
| | | 1.0 | 9810 ± 1090 | 57 ± 41 |
| | | 1.7 | 25552 ± 2744 | 227 ± 113 |
| | R- | 0.7 | 1785 ± 185 | 28 ± 9 |
| | | 1.0 | 9029 ± 1380 | 100 ± 30 |
| | | 1.7 | 16602 ± 1667 | 190 ± 48 |
| Winter 2 | R+ | 0.3 | 1400 ± 466 | 12 ± 5 |
| | | 1.0 | 8615 ± 2372 | 104 ± 38 |
| | UV- | 0.3 | 610 ± 440 | 3 ± 1 |
| | | 1.0 | 6851 ± 1841 | 50 ± 27 |
| | R- | 0.3 | 1499 ± 206 | 7 ± 4 |
| | | 1.0 | 10809 ± 1319 | 66 ± 21 |

R+: full radiation. UV-: UV blocked. R-: UV, blue and green light blocked.

Table S4: ANOVA results of β-glucosidase and phenol-oxidase accumulated activities for the light attenuation treatment by species and year.

| Species / Group | Year | Degrees of freedom | $R^2$ | *p* |
|---|---|---|---|---|
| β-glucosidase | | | | |
| *P. stuckertii* | | | | |
| Winter Group 1 | 0.3 | 12 | 0.3 | 0.1 |
| | 1.0 | 12 | 0.2 | 0.3 |
| | 1.3 | 12 | 0.06 | 0.7 |
| | 2.1 | 12 | 0.2 | 0.2 |
| Spring Group | 0.7 | 12 | 0.4 | 0.04 * |
| | 1.0 | 12 | 0.2 | 0.4 |
| | 1.7 | 12 | 0.1 | 0.5 |
| Winter Group 2 | 0.3 | 12 | 0.06 | 0.7 |
| | 1.0 | 12 | 0.06 | 0.7 |
| *D. hieronymi* | | | | |
| Winter Group 1 | 0.3 | 12 | 0.06 | 0.7 |
| | 1.0 | 12 | 0.4 | 0.2 |
| | 1.3 | 12 | 0.07 | 0.6 |
| | 2.1 | 12 | 0.5 | 0.02 * |
| Spring Group | 0.7 | 12 | 0.06 | 0.7 |
| | 1.0 | 12 | 0.2 | 0.3 |
| | 1.7 | 12 | 0.5 | 0.01 * |
| Winter Group 2 | 0.3 | 11 | 0.2 | 0.3 |
| | 1.0 | 11 | 0.2 | 0.3 |
| Phenol-oxidase | | | | |
| *P. stuckertii* | | | | |
| Winter Group 1 | 1.0 | 10 | 0.006 | 0.97 |
| | 1.3 | 12 | 0.03 | 0.8 |
| | 2.1 | 11 | 0.2 | 0.3 |
| Spring Group | 0.7 | 9 | 0.2 | 0.4 |
| | 1.0 | 9 | 0.4 | 0.1 |
| | 1.7 | 9 | 0.05 | 0.7 |
| Winter Group 2 | 0.3 | 12 | 0.05 | 0.7 |
| | 1.0 | 12 | 0.06 | 0.7 |
| *D. hieronymi* | | | | |
| Winter Group 1 | 1.0 | 12 | 0.009 | 0.8 |
| | 1.3 | 12 | 0.009 | 0.9 |
| | 2.1 | 11 | 0.1 | 0.5 |

| | | | | |
|---|---|---|---|---|
| Spring Group | 0.7 | 10 | 0.5 | 0.03 * |
| | 1.0 | 10 | 0.1 | 0.5 |
| | 1.7 | 10 | 0.02 | 0.9 |
| Winter Group 2 | 0.3 | 11 | 0.2 | 0.4 |
| | 1.0 | 11 | 0.1 | 0.4 |

Asterisk denote significant results: * $p<0.05$.

Table S5: ANOVA results for decomposition rates by species and time.

| *Species / Group* | *Year* | Degrees of freedom | $R^2$ | p |
|---|---|---|---|---|
| *P. stuckertii* | | | | |
| Winter Group 1 | 1 | 12 | 0.3 | 0.09 |
| | 2 | 11 | 0.2 | 0.4 |
| Spring | 1 | 12 | 0.1 | 0.5 |
| | 1.7 | 9 | 0.6 | 0.03 * |
| *D. hieronymi* | | | | |
| Winter Group 1 | 1 | 11 | 0.6 | 0.006 ** |
| | 2 | 8 | 0.9 | 0.0003 *** |
| Spring Group | 1 | 12 | 0.8 | 0.0001 *** |
| | 1.7 | 12 | 0.8 | < 0.0001 *** |

Asterisks denote significant results: *** $p<0.001$, ** $p<0.01$, * $p<0.05$.

Table S6: Initial litter traits (± SE) and ANOVA results by species.

| Trait | *P. stuckertii* | *D. hieronymi* | df | $R^2$ | *p* |
|---|---|---|---|---|---|
| A305 | 0.38 ± 0.028 | 0.49 ± 0.019 | 7 | 0.7 | 0.008 ** |
| Celullose (%) | 43.34 ± 0.543 | 37.86 ± 0.37 | 8 | 0.9 | < 0.0001 *** |
| Hemicellulose (%) | 48.45 ± 0.82 | 52.73 ± 0.68 | 8 | 0.7 | 0.004 ** |
| Lignin (%) | 8.19 ± 0.38 | 9.41 ± 0.77 | 8 | 0.2 | 0.2 |
| Saccharification (mg $g^{-1}$) | 172.6 ± 11.84 | 199.71 ± 5.38 | 8 | 0.4 | 0.07 |
| Solluble carbohydrates (%) | 5.14 ± 0.70 | 5.22 ± 0.98 | 8 | 0.0006 | 0.9 |
| Total polyphenols (mg $g^{-1}$) | 0.079 ± 0.006 | 0.066 ± 0.007 | 8 | 0.2 | 0.2 |
| Total sugars (mg $g^{-1}$) | 0.12 ± 0.015 | 0.14 ± 0.002 | 7 | 0.2 | 0.2 |
| LMA (mg $mm^{-2}$) | 0.32 ± 0.018 | 0.112 ± 0.010 | 8 | 0.9 | < 0.0001 *** |
| Potential leaching (%) | 0.68 ± 0.22 | 0.085 ± 0.085 | 8 | 0.4 | 0.03 * |
| Toughness (N $mm^{-1}$) | 4.46 ± 0.42 | 1.14 ± 0.17 | 7 | 0.96 | < 0.0001 *** |
| Water adsorption capacity (%) | 81.61 ± 5.52 | 64.88 ± 3.57 | 8 | 0.4 | 0.03 * |

df: degrees of freedom. A305: sunscreens with absorptance at 305 nm. LMA: leaf mass per area. Asterisks denote significant results: *** $p<0.001$, ** $p<0.01$, * $p<0.05$.

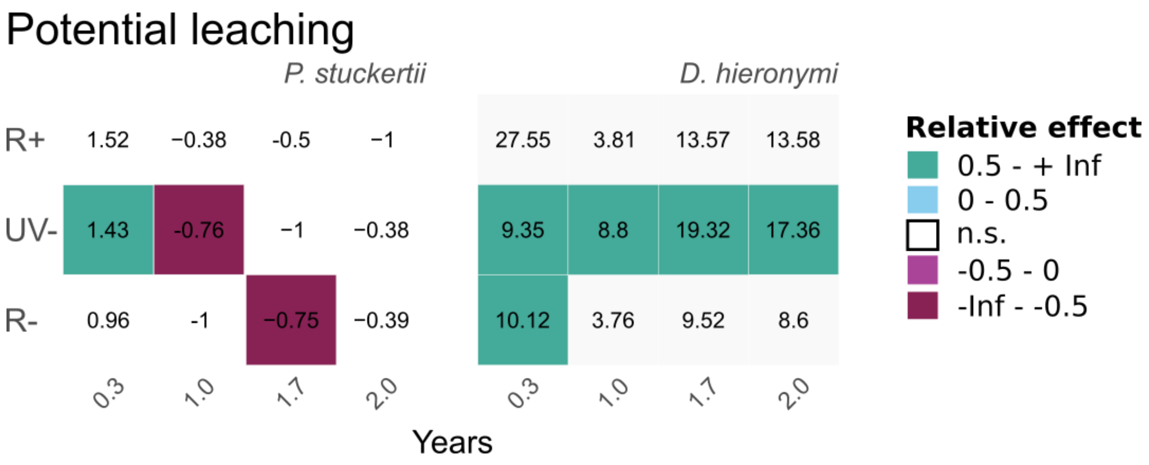


Figure S3: Relative effect of field decomposition on potential leaching (initial weight lost after soaking, %) compared to initial values for both species under each filter. Colored squares denote significant differences for a t-test (*p<0.05*). Colors represent direction of change and intensity: blue shades represent increases in trait values and pink shades represent decreases, while lighter shades represent low-moderate changes (0 – ±0.5) and darker shades represent bigger changes (±0.5 – infinite).
